# Time-varying Dynamic Network Model For Dynamic Resting State Functional Connectivity in fMRI and MEG imaging

**DOI:** 10.1101/2021.04.01.438060

**Authors:** Fei Jiang, Huaqing Jin, Yijing Gao, Xihe Xie, Jennifer Cummings, Ashish Raj, Srikantan Nagarajan

## Abstract

Dynamic resting state functional connectivity (RSFC) characterizes fluctuations that occurs over time in functional brain networks. Existing methods to extract dynamic RSFCs, such as sliding-window and clustering methods, have various limitations due to their inherent non-adaptive nature and high-dimensionality including an inability to reconstruct brain signals, insufficiency of data for reliable estimation, insensitivity to rapid changes in dynamics, and a lack of generalizability across multimodal functional imaging datasets. To overcome these deficiencies, we develop a novel and unifying time-varying dynamic network (TVDN) framework for examining dynamic resting state functional connectivity. TVDN includes a generative model that describes the relation between low-dimensional dynamic RSFC and the brain signals, and an inference algorithm that automatically and adaptively learns to detect dynamic state transitions in data and a low-dimensional manifold of dynamic RSFC. TVDN is generalizable to handle multimodal functional neuroimaging data (fMRI and MEG/EEG). The resulting estimated low-dimensional dynamic RSFCs manifold directly links to the frequency content of brain signals. Hence we can evaluate TVDN performance by examining whether learnt features can reconstruct observed brain signals. We conduct comprehensive simulations to evaluate TVDN under hypothetical settings. We then demonstrate the application of TVDN with real fMRI and MEG data, and compare the results with existing benchmarks. Results demonstrate that TVDN is able to correctly capture the dynamics of brain activity and more robustly detect brain state switching both in resting state fMRI and MEG data.

## 1 Introduction

The human brain can be described as highly dynamic functional networks constructed from a fixed structural structural network whose fluctuations over time form the basis for complex cognitive functions and consciousness (Bassett et al., 2011; Deco & Jirsa, 2012; Shine et al., 2015). This view of brain function highlights the importance of time sensitive descriptions of brain network activity in understanding the functional relevance of alterations in network structure that may underlie different behavioral states and conditions(Varela et al., 2001). Recent experiments using fMRI data have demonstrated that global brain signals transition between states of high and low connectivity strength over time (Zalesky et al., 2014) and these fluctuations are related to coordinated patterns of network topology (Betzel et al., 2016). Studies suggest that dynamic fluctuations in network structure relate to that in cognitive function (Shine et al., 2015). Therefore, analyses of functional neuroimaging data to examine time-varying reconfiguration of global network structure may provide a unique opportunity to gain insights into the dynamics of functional brain networks, their association with behavioral states, and their alterations in disease and therapeutic interventions.

To appropriately describe synchronous temporal fluctuations in neuroimaging data, many data driven approaches have been used, especially with resting state functional connectivity (RSFC) which describes how brain activity is correlated across regions when an explicit task is not being performed. Many studies have shown that this functional connectivity provides a powerful and informative framework for exploring brain organization (Bullmore & Sporns, 2009; Greicius, 2008; Shine et al., 2015). RSFC studies have been described both for blood-oxygen level-dependent (BOLD) data measured with functional magnetic resonance imaging (fMRI) (Biswal et al., 1997; Calhoun et al., 2001; Greicius et al., 2003) and for faster time scale neural oscillatory network changes measured with magnetoencephalography (MEG) (Englot et al., 2015; Ranasinghe et al., 2017) or electroencephalography (EEG) imaging (Brookes et al., 2011; Dominguez et al., 2013; Hohlefeld et al., 2013).

Approaches for RSFC analyses include seed-based correlations (Lv et al., 2018), independent component analysis (Beckmann et al., 2005) and dynamic mode decomposition (Brunton et al., 2016; Kutz et al., 2016). Recent work has also focused on recovering the static RSFC from underlying structural connectivity via graph methods like network diffusion model (Abdelnour et al., 2014) and algebraic spectral graph expansions (Abdelnour et al., 2018; Becker et al., 2018; Meier et al., 2016; Tewarie et al., 2020). However, few statistical methods have been proposed for the accurate estimation of dynamic changes in functional network architecture (Shine et al., 2015). To date, most existing statistical techniques for RSFC have assumed that the functional connectivity structure is stationary over a dataset, which is in direct contrast to emerging data that suggests that the strength of connectivity between regions is variable over time. Therefore, the development of statistical methods that enable exploration of dynamic changes in functional connectivity is currently of great importance to the neuroscience community.

The extension of current techniques to capture the dynamic changes in RSFC during the scan period is a lively yet evolving topic. It is well known that the brain at rest is in fact quite dynamic, with RSFC capable of changing over a matter of seconds to minutes (Hutchison et al., 2013). This time varying pattern, namely dynamic resting state functional connectivity, has been shown to constitute novel imaging biomarkers for identifying neurological dysfunctions such as schizophrenia, autism and various forms of dementia (Damaraju et al., 2014; Filippi et al., 2019; Ma et al., 2014; Mash et al., 2019; Rashid et al., 2016, 2014; Schumacher et al., 2019). For instance, it can identify early mild cognitive impairment for dementia (Wee et al., 2016) and distinguishes Alzheimer’s Disease (AD) patients from healthy controls (Schumacher et al., 2019). Thus, the dynamic component of RSFC may serve as additional biomarker of neurological disorders - a key motivation of current work.

Currently, the most common approach to extract dynamic RSFC relies on the sliding-window method, which generally consists of two steps: (1) divide signals into segments of equal duration; (2) implement the traditional seed based method (Biswal et al., 1995; Fox et al., 2005), independent component analysis (Allen et al., 2014; Calhoun et al., 2001; van de Ven et al., 2004), or the dynamic mode decomposition method (Brunton et al., 2016; Kutz et al., 2016) on the segments sequentially.

While the sliding-window is practically attractive since it enables the use of earlier static methods in the dynamic context, it presents several limitations and trade-offs, which we will discuss in Section 3.1. We present a unified solution for extracting dynamic RSFCs from both fMRI and MEG data, which directly addresses these limitations. We call this method the time-varying dynamic network (TVDN) framework. We develop a novel automatic and provably statistically optimal inference algorithm based on the TVDN model to infer the dynamics that underlie the model. We extract the stationary spatial features and detect the dynamic brain state switches adaptively. The algorithm is able to divide the brain signals into uneven segments, each of which contains brain activities in a stationary mental state.

Once the parameters have been successfully inferred, the entire spatio-temporal noise-free imaging signal can be reconstructed through a high dimensional linear forward model - a feature that is rarely available in current methods. The algorithm involves a few tuning or hyper-parameters, which are automatically selected to minimize the uncertainties of number of switches across independent samples. We expect that the presented TVDN framework will prove effective in robustly generating dynamic RSFC features that will serve as useful biomarkers of neurological and neurodegenerative diseases.

## 2 Results

### 2.1 A new biologically constrained model of dynamic functional connectivity

Current methods for dynamic functional connectivity (FC) analysis do not account for biological constraints or biophysically realistic models of brain activity and state switches. This represents a lost opportunity to overcome some of the limitations noted in Section 3.1. Here we propose a novel model of dynamic FC that relies on discrete and discontinuous “state changes” in brain activity. Indeed, there is mounting evidence that the brain’s dynamics results from its cycling through a number of micro-states separated by brain state switches, such that while the FC between state switches may be considered stationary, their transitions are subject to discontinuous, abrupt or non-smooth events. In addition to being more biologically realistic, this approach allows us to benefit from several constraints, especially the concept that the spatial features of brain activity might be stationary, while the coupling between these stationary structures might be temporally dynamic. For instance, the spatial structures may arise from the underlying structural connectivity, while the temporal parameters describe the dynamic switching between brain networks over time. Therefore, while the spatial structure of the FC patterns are considered stationary due to the linkage with the structure of the brain, how these spatial features work together is allowed to vary over time. It is further possible to constrain the dynamics of the temporal parameters. Rather than randomly or continuously traversing through the latent state-space, RSFCs most likely undergoes discrete and discontinuous shifts, resulting in the concept of “microstates”. Hence we impose piece-wise constancy to these temporally changing coefficients. We show that using these powerful constraints, it is possible to overcome the trade-offs and limitations currently pertinent to dynamic RSFC analysis.

Let **X**(*t*) be a vector of the brain signals at time *t* from all brain regions of interest, and **X**’(*t*) be the corresponding signal increment at time *t*. To incorporate the static spatial and dynamic temporal features, we model the increment of signal at a specific imaging acquisition time *t* as

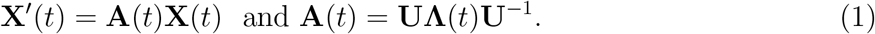

Here **U** represents the static spatial feature while **Λ**(*t*) represents the piece-wise dynamic temporal feature. The rank of **A**(*t*) represents the number of brain states. This model suggests that at each stationary state in between two brain switches, the brain signal follows

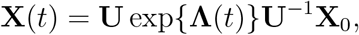

where **X**_0_ is the initial signal strength. We discuss the motivation of the model in Section 4 in more detail.

Let **Y**_*j*_ be the observed signal sampled at the *j*th time such that **Y**_*j*_ = **X**(*t_j_*) + ϵ_*j*_, and let ϵ_*j*_ be random noise. Based on a sequence of **Y**_*j*_, *j* = 1,…,*n* across the acquisition times, TVDN uses a kernel regression approach to estimate **U** and a brain state switch detection method to capture the changes in **Λ**(*t*). Figure 1 shows a flowchart of the estimation procedure. The purple ovals represent TVDN inputs, and red ovals represent TVDN outputs. The blue rectangles represent the building blocks of TVDN, which we discuss in detail in Section 4.3.3. In the following results we use 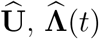 and 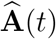 to denote the esti-mators for **U, Λ** and **A**(*t*), respectively. From (4) in Appendix, 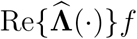 is the estimator of growth/decay constant and 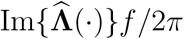 is the estimator for the signal frequency in Hertz, where 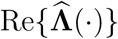 and 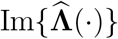 are the real and imaginary part of 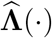 and *f* is the signal sampling frequency.

**Figure 1.**
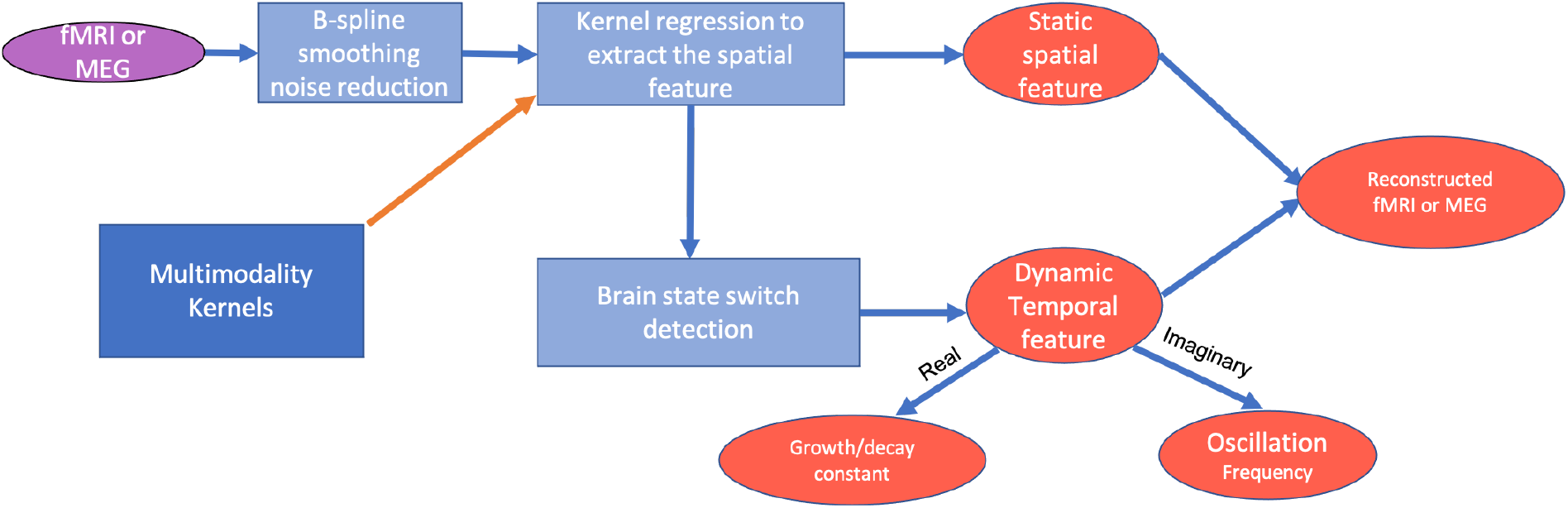
TVDN pipline. The purple ovals represent TVDN inputs, and red ovals represent TVDN outputs. The blue rectangules represent the building blocks of TVDN. Two multimodality kernel examples are provided and will be discussed in Section 4.3.3.

We also implemented three sliding-window based approaches for the comparison. Detailed implementation procedure is discussed in Section 4.4.

### 2.2 Simulation study

We implement TVDN on brain signals simulated from Model (1). We also implemented three sliding-window based approaches for the comparison on the same data. Detailed simulation procedure and the implementation of the sliding window methods are discussed in Section 4.5 and 4.4, respectively.

We plot the estimated switches in Figure 2 (a). The result shows that TVDN captures the brain state switch accurately. To illustrate the estimation results, we reconstruct the data by using estimated spatial and temporal features. We show the mean of the estimators at selected brain regions and the 95% empirical confidence interval, that is 5% and 97.5% quantiles of the estimators over 100 simulations in Figure 2 (b). Figure 2 (b) shows that TVDN recovers the original noiseless sequence and the confidence intervals cover the true signals.

**Figure 2:**
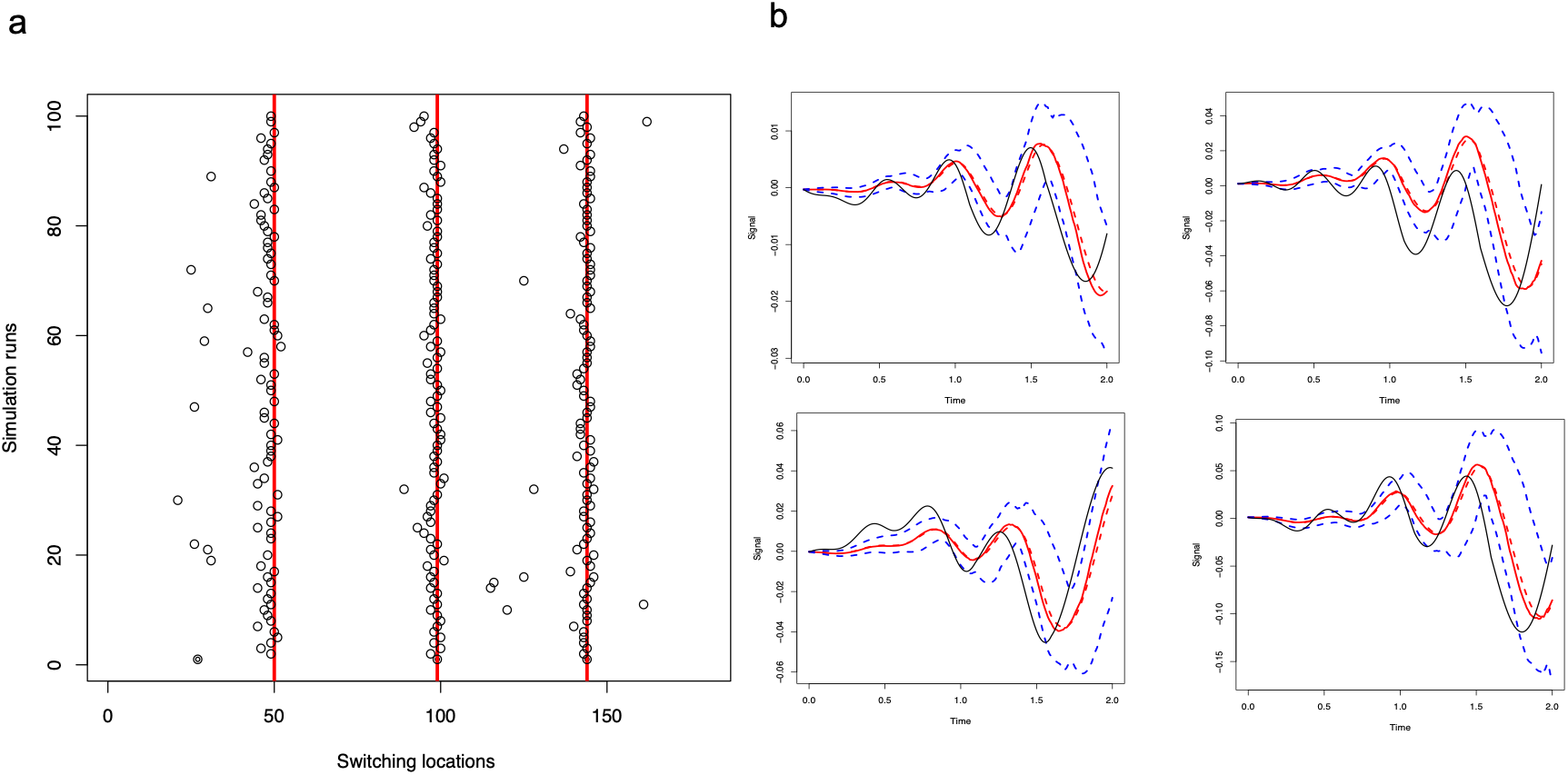
The simulation results with three switches. TVDN detect the true brain state switches and can reconstruct the true signal. The sliding-window methods are sensitive to the window size. (a) Switch times. Red lines are the true switch times and the dots are the estimated locations. (b) The black lines are the true **X**(*t*) at four selected regions. The red solid and dash curves are the mean and median of the estimators and above and below blue curves are the 95% empirical confidence intervals. The figures from left to right represent the results of the estimators whose mean squared errors fall at the 0%, 25%, 50% and 75% quantiles of the mean squared errors across all simulations.

We also plot the resulting switch locations from the sliding-window methods in Figure 9 in Section 4.5. Figure 9 (c) shows that none of the three methods correctly identify the switches. In addition, the sliding-window based methods are sensitive to the window size changes, which leads to substantial different results when varying the window sizes.

### 2.3 TVDN results for resting state fMRI data

We present the TVDN detection results from one fMRI sequence in Figure 3 (a). We also obtain the growth/decay constant 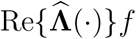 and the signal frequency as 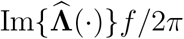, where *f* = 0.5 Hz is the sampling frequency of the fMRI signal. It can be seen from Figure 3 (b) that the resting state fMRI brain signals are active in the frequency range between 0.001 and 0.007 HZ. We calculated the Pearson correlation between the weighted spatial features, that is, column sum of 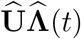, and the seven canonical networks from Yeo et al. (2011)‘s independent component analysis. As shown in Figure 3 (c), the subject’s weighted spatial features have the strongest correlation with the limbic network in the first segment (0.38), with the ventral attention network in the second (0.443), third (0.415), sixth (0.432) and eighth (0.32) segments, with the dorsal attention network (0.24) in the fourth segment. This changing correlation pattern is indicative of brain state switches over time, demonstrating that different functional networks are operational at various times. In order to visualize the changing spatial patterns, we plotted the weighted spatial features across the segments on the brain surface in Figure 3 (d). The results again illustrate that the spatial pattern reflecting brain state switches among frontal, parietal and occipital lobes over time.

**Figure 3:**
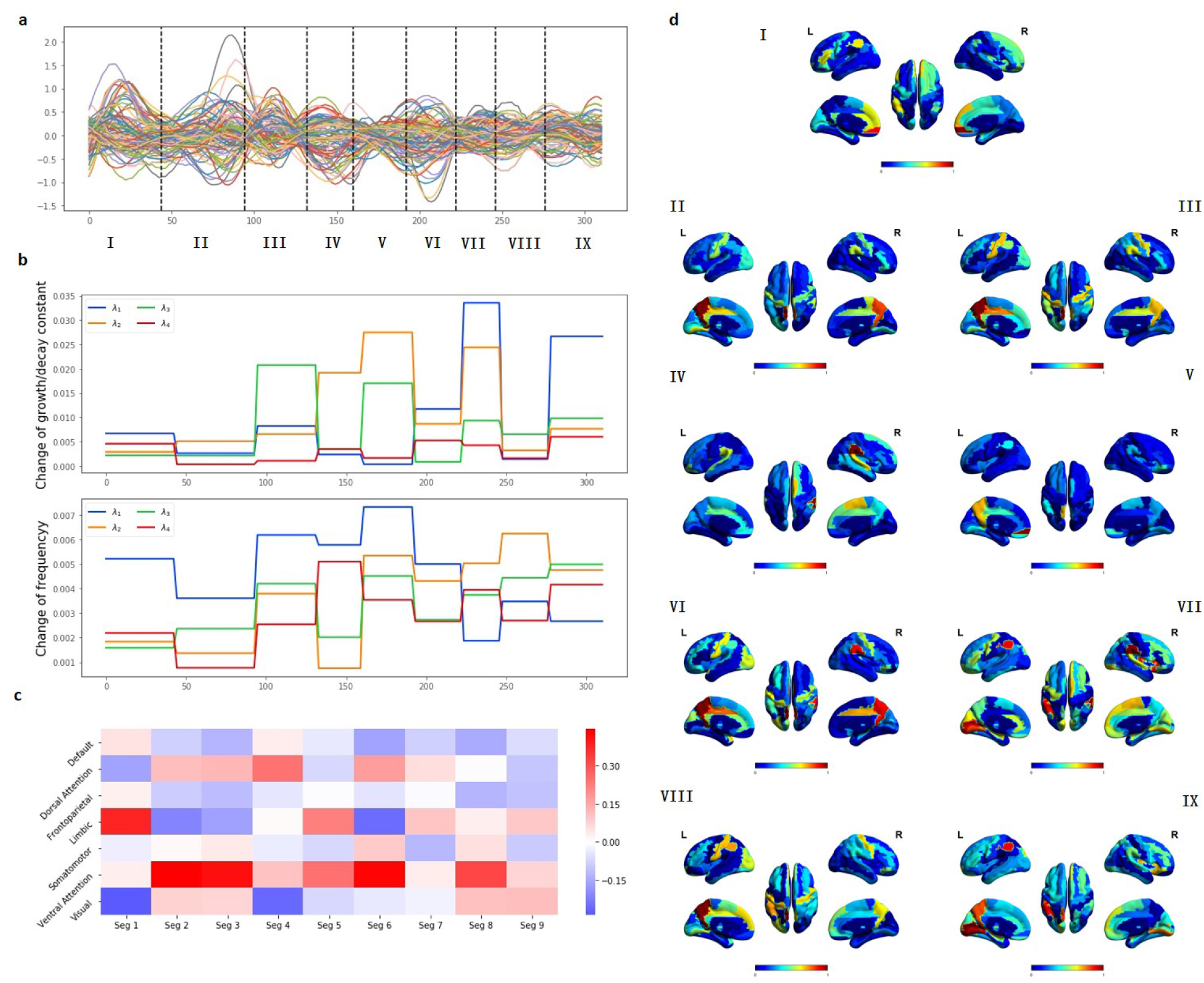
The results from the first fMRI dataset. (a) The the real sequences with switch locations (black dash lines) detected by TVDN. (b) Changes of growth/decay constant 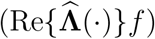, changes of the frequencies 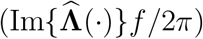. (c) The Pearson correlation be-tween the weighted spatial features and the seven canonical networks. (d) The weighted spatial features across different segments detected by TVDN.

We also plot, in Figure 11 in Appendix, the pair-wise connectivity measure in each segment, defined by exp(-||**x**_1_ — **x**_2_||_2_), where **x**_1_, **x**_2_ represent signal sequences from two brain regions. Figure 11 shows that the connectivity increases gradually over time. For comparison we show analogous results from TVDOR, TVPCA and TVDMD methods with different window sizes in Figure 12 in the Appendix. The latter results suggest that these existing sliding-window methods are sensitive to tuning parameters and do not give coherent switch times when different window sizes are selected. Another representative example similar to the above is given in Figure 13; its connectivity measures in Figure 14 and the results from competing methods 15 in Appendix.

### 2.4 TVDN results for resting state MEG data

We evaluated TVDN on resting state MEG data, where we consider series of de-trended MEG source signals with *d* = 68 ROIs. Note that for MEG data, we did not filter the source signals because the high time resolution MEG data contain clear fluctuation trends that are not overwhelmed by the noise.

We obtain the detection results as shown in Figure 4. We also obtain the growth/decay constant 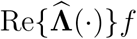 and the signal frequency as 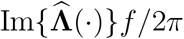, where *f* = 60 is the sampling frequency for the MEG signal. The results in Figure 4 (a) show that there are seven switches in the signal. In addition, the brain is active in the frequency range between 0 to 6 Hz as shown in Figure 4 (b). We also plot the correlation between the weighted spatial features and the seven canonical network in Figure 4 (c). It can be seen that the subject’s weighted spatial features have the strongest correlation with the visual network in the first segment (0.31), with the dorsal attention network in the second (0.32), third (0.45), fifth (0.28) and sixth (0.44) segments, with limbic network (0.29) in the seventh segment, and with frontopartietal network (0.62) in the eighth segment. These correlation are larger than those from the resting state fMRI (Figure 3). Finally, we view the weighted spatial features across the segments in Figure 4 (d). The results illustrate the characteristic brain state activation patterns switching among parietal, frontal, temporal and occipital lobes over time. We also plot, in Figure 16 in Appendix, the pair-wise connectivity measure in each segment, which shows that the connectivity increases from the first to the third segment, decreases from the fourth to the sixth segment, and increases again until the end of the time. We further show the results from the existing sliding-window methods in Figure 17 in the Appendix, which demonstrates that the sliding-window methods are sensitive to window size selection. Another representative example is given in Figure 18, the corresponding connectivity measures in Figure 19 and results from competing methods in Figure 20 in Appendix.

**Figure 4:**
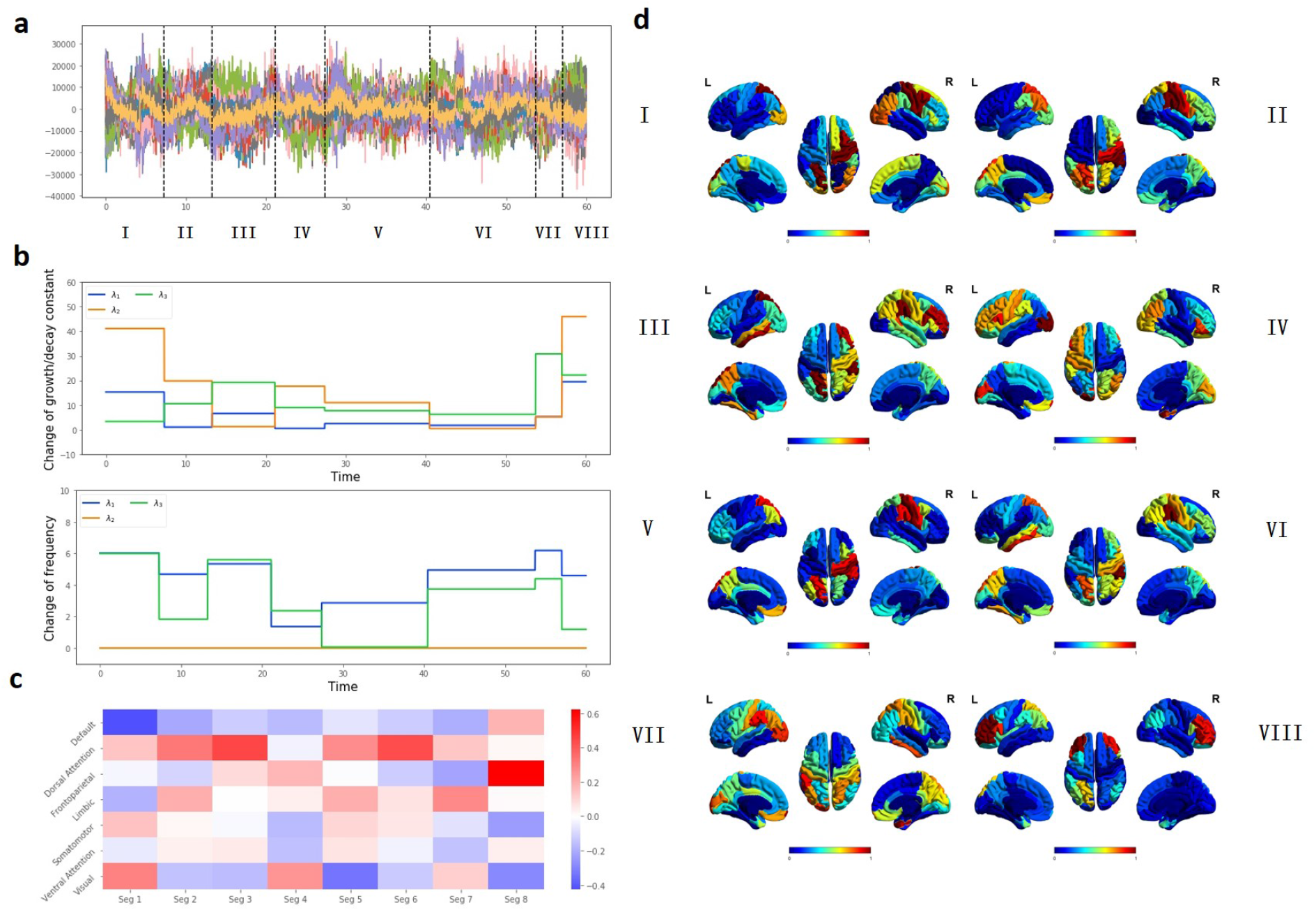
The results from the first resting state MEG dataset. (a) The the real sequences with switch locations detected by TVDN. The dash line is the detected brain state switches. (b) Changes of growth/decay constant 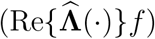, changes of the frequencies 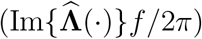. (c) The Pearson correlation between the weighted spatial features and the seven canonical networks. (d) The weighted spatial features across different segments detected by TVDN with eye-opening-closing labels.

### 2.5 TVDN results for task based MEG data

To validate the accuracy of brain state switch detection, we evaluate TVDN on MEG recordings during a simple eyes-open to eyes-close task-switching experiment, where six eye close and open tasks blocks were performed within one minute and the switch times were manually labeled. In Figure 5, we show the detection results based on the MEG data from two subjects. Clearly, the switch locations from TVDN are very close to the manually labeled ones, which suggest TVDN can correctly identify the brain state switch times. Take the first sample as an example, we obtain the growth/decay constant and the signal frequency with *f* = 120 be the sampling frequency for the two task based MEG signals. The brain is active in the frequency between 0 to 12 as shown in Figure 21 (a) in Appendix, which is higher than that from the resting state MEG. Furthermore, we obtain the band passed signals in alpha band (8-12 HZ), and re-estimated the **U** and **Λ** based on the filtered the signals. We then calculate the Pearson correlation between the re-estimated weighted spatial features and the seven canonical networks. As shown in Figure 21 (b) in Appendix, although the correlations with the visual network change over time, the switch patterns do not exactly follow the eyes-open and eyes-close states. This implies there are micro-state changes that are unrelated to the visual network during the data acquisition period. Moreover, the brain views of the re-estimated weighted spatial features in Figure 21 (c) illustrate that the brain state in alpha band switches in between inferior parietal and supra marginal in the parietal lobe in most of the segment, while it switches to occipital lobe at the end of the time. We also plot in Figure 22 in Appendix the pair-wise connectivity measure in each segment base on the unfiltered signal, which shows that the connectivity decreases from the first to the fourth segment, increases from the fourth to the fifth segment, and decreases again to the end of the time. We further show the results from the TVCOR, TVPCA, and TVDMD methods in Figure 23 in Appendix, which suggests none of the methods provide robust result across the selected window sizes. Finally, base on the second sample, we show the growth decay constants, signal frequency, correlations with the canonical networks and brain views in Figure 24, the connectivity measures in Figure 22 and the results from competing methods in Figure 26 in Appendix.

**Figure 5:**
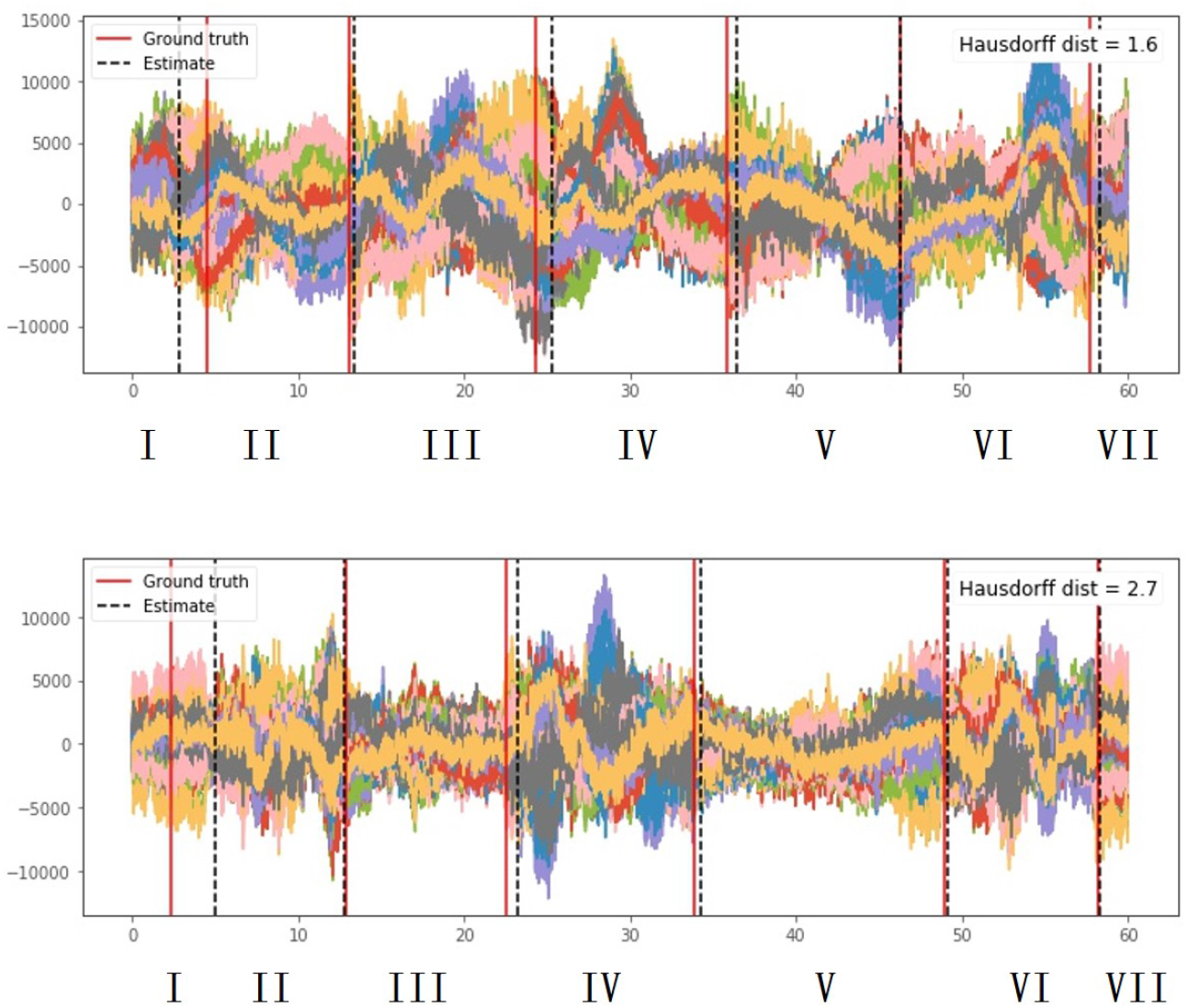
TVDN captures the task-switching dynamics in two eyes-open to eyes-close task-switching MEG records. The the real sequences with switch locations detected by TVDN. The black dashed lines are the detected brain state switches. The red solid lines are the manually labeled switch times.

### 2.6 Comparison to benchmark methods

We implemented TVDN on 103 fMRI datasets. The distribution of the number of switches and ranks are displayed in Figure 6 (a) and (b), respectively, which show around 50% samples have eight switches and over 65% samples have seven distinct brain states (ranks) in the resting state.

We further evaluate the correlations between TVDN spatial features with the seven canonical networks from Yeo et al. (2011)‘s independent component analysis under selected *κ* and *r*. We extract the spatial features as the modulus of the first *r* columns of 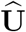 from each subject, and project them to [0,1] interval. We also implemented the TVPCA, TVDMD methods to obtain the corresponding principle components and dynamic modes from each segment as the spatial features, and calculated their correlations with the canonical net-works. We plot the distributions of the maximum correlations between the canonical networks and the spatial features from TVDN, TVPCA and TVDMD across 103 samples in Figure 6 (c). It can be seen that although TVDN has far fewer spatial features compared with TVPCA and TVDMD (each subject has only *r* spatial features), the distribution of the maximum correlation is similar with those from TVPCA and TVDMD. In addition, we plot the prediction errors versus the number of switches from TVDN and TVDMD in Figure 6 (d). To obtain the prediction error, for each segment in between two consecutive switch points, we use the first half of the fMRI records as the training data to estimate **A**(*t*) in the segments. Then we use the rest of the signals as the testing data, and calculate the related prediction errors defined as

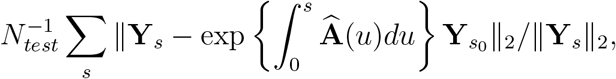

where **Y**_*s*_ is the sth observed signals, **Y**_*s*0_ is first signal in the testing sample, *N_test_* is the total number of testing sample (half of the signal length) and the summation is over the test signals. We average the corresponding prediction errors across the segments and individuals for the TVDN and TVDMD methods. For the TVDN method, we further construct the 95% confidence bands of the prediction errors as 2.5 (lower) and 97.5 (upper) quantiles of the erros in the 103 study samples. We do not show the 95% confidence band from TVDMD because it almost covers the entire plotted area. Figure 6 (d) shows that TVDN has smaller prediction error than TVDMD, especially when the number of switches is larger than four. It also suggests that when the number of switches is small, each segment contains sufficient samples to recover the large number of parameters in TVDMD. Therefore, when there are less than four switches, TVDN and TVDMD perform equally well in prediction as the confidence band cover both curves. However, when the number of switches is moderately large, the sample in each segment is no longer enough to provide accurate estimations for TVDMD parameters. Therefore, the more parsimonious TVDN method yields substantially smaller prediction errors than the TVDMD method.

**Figure 6:**
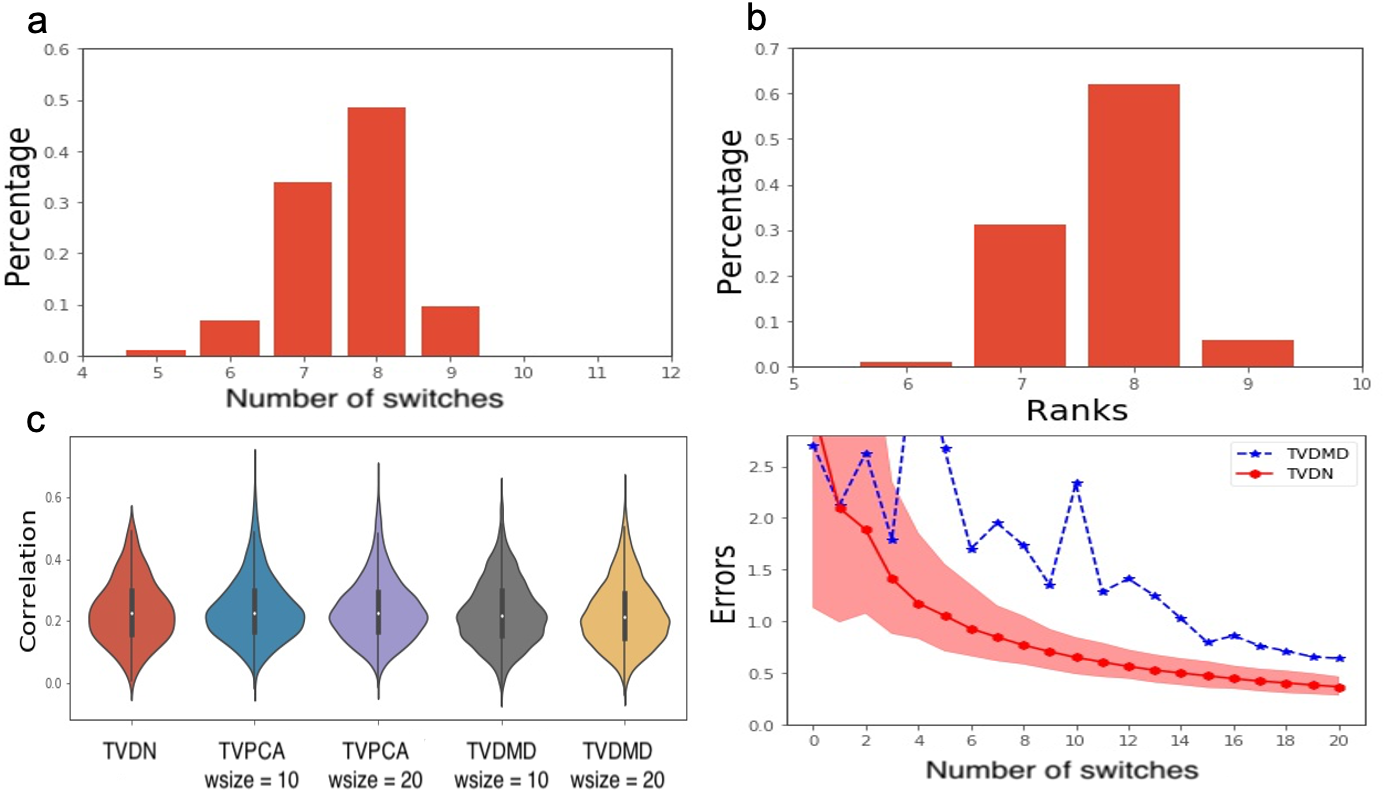
The distribution of the spatial features are similar from different approaches (c) and the related prediction error from TVDN is smaller compare with those from TVDMD, the only existing method that allows for the reconstruction of the signals (d). (a) The distribution of the number of switches across samples. (b) The distribution of the brain state across samples. (c) The distributions of the maximum correlation between the spatial features from TVPCA, TVDMD, and TVDN methods with the canonical networks. (d) The average related prediction errors from the TVDN and TVDMD methods. The shaded area is the 95% confidence band from the TVDN method. The 95% confidence band from TVDMD covers the entire plotted area, which we do not show.

To illustrate the robustness of TVDN, in Figure 7, we plot the distribution of the number of switches from TVDN and the sliding-window methods when different kernel bandwidths and window sizes are selected, respectively. Note that the kernel bandwidth in TVDN serves the same function as the window sizes in the sliding-window methods. For each window size, we adjust the kernel bandwidth so that the lower 2.5% and upper 97.5% of the Gaussian kernel correspond to the left and right endpoints of the window, respectively. It can be seen that TVCOR, TVPCA and TVDMD are sensitive to the window size selection - the larger the window size, the smaller the number of detected brain switches. In contrast, TVDN is robust to the kernel bandwidth selection, with only small shifts of the distribution center with increasing kernel bandwidth.

**Figure 7:**
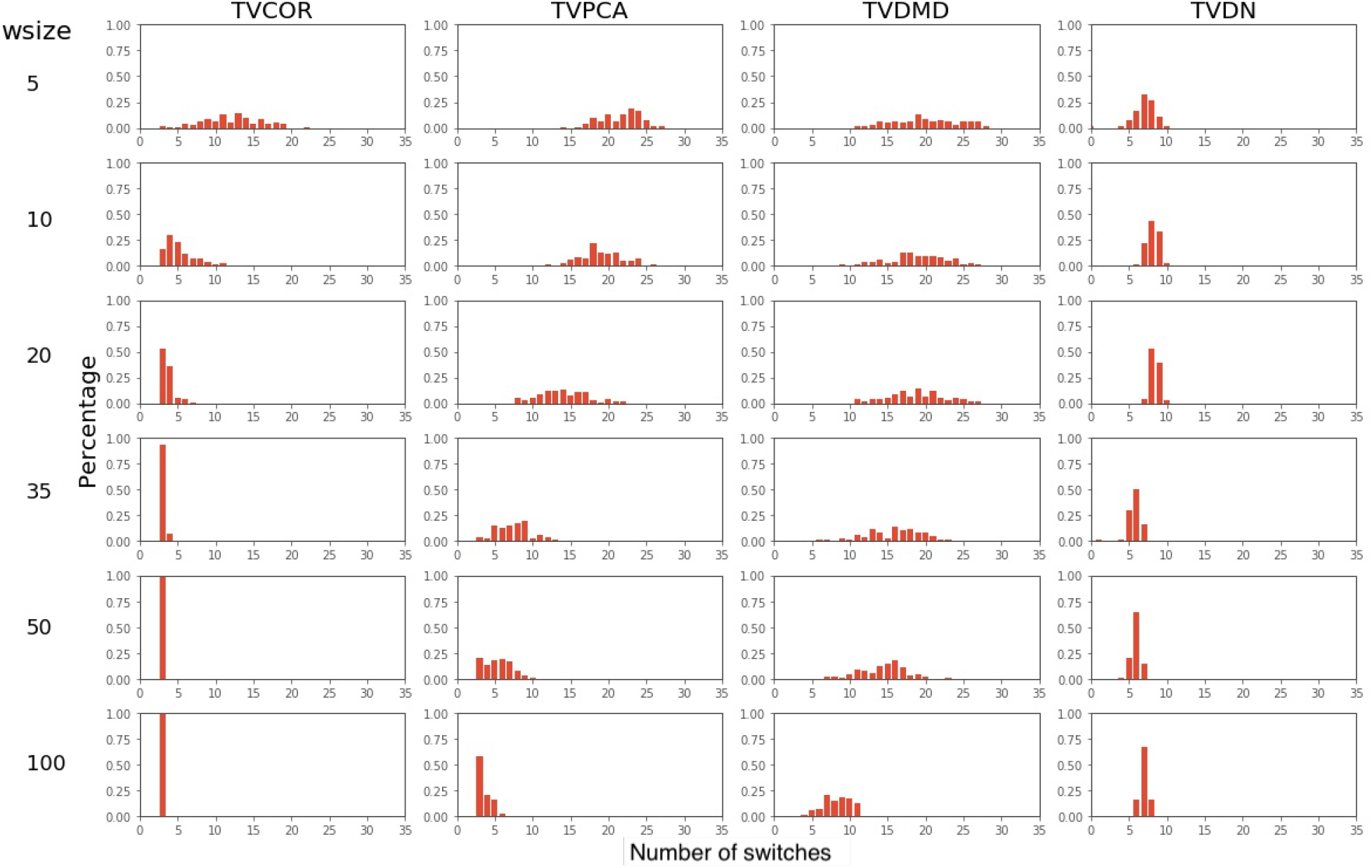
TVDN’s brain state switch detection is robust to the kernel bandwidth selection but the sliding-window methods are sensitive to window size selection. The distribution of the switch points when different window sizes (wsize) are chosen for the sliding-window methods and different kernel bandwidths are chosen for TVDN. The kernel bandwidths are adjusted so that the lower 2.5% and upper 97.5% of the Gaussian kernel correspond to the left and right endpoints of the window, respectively.

## 3 Discussion and Conclusion

We proposed a novel biologically-constrained model of brain state evolution during restingstate functional recording, called the TVDN model. We presented an optimal algorithm to infer the model’s parameters and to extract the spatial and temporal features from resting state brain signals. The method relies on the assumption that while the spatial signatures of RSFC, given by the eigenvectors of the forward model, are static, the evolution of temporal features, given by the eigenvalues, is dynamic within the recording duration. We developed an eigenvector estimation technique to extract consistent spatial features across signal acquisition times. In addition, we proposed a dynamic programming-based algorithm to detect temporal switches adaptively based on the signal oscillation patterns, under the biologically-inspired assumption that state transitions are abrupt rather than smooth in time. Using the inferred spatial and temporal features, we can reconstruct the underlying mean signals that generate the noisy observations. This may be considered a model-based smoothing operation, with several potential applications. Thus, our method is a legitimate generative model of dynamic functional activity in the brain. In addition, the ability to reconstruct noiseless signals gives the algorithm an opportunity to tune its parameters using a reconstruction error metric to be minimized.

We evaluated the method on thorough simulated data, followed by a rigorous characterization of its performances on empirical fMRI and MEG data from the BIL laboratory at UCSF. The simulation study shows that TVDN captures the true brain switch locations and is able to recover the true signal that generates the observed ones. In the empirical study, for comparison we implemented several competing technques, including TVCOR, TVPCA and TVDMD methods. Compared with competing methods, TVDN produces smaller set of spatial features but their correlations with the seven canonical networks have the same distributions as those from the TVCOR, TVPCA and TVDMD methods. This suggests the smaller set of spatial features from TVDN is sufficient to explain the brain connection patterns. Furthermore, TVDN provides more robust temporal features, which are adaptive to the signals and noises from different data and is insensitive to the tuning parameters, such as kernel bandwidth. In addition, the evaluation on the eye-opening-closing task data shows that TVDN captures the brain state switches accurately. More importantly, TVDN has significantly smaller prediction errors than TVDMD does when predicting “future” activity in the same segment. Last but not least, the resulting temporal features include instantaneous estimates of the active oscillation frequency of functional activity, thus imparting the method with attributes of a model-based alternative to conventional time-frequency analysis.

The ultimate solution to improving the estimation accuracy on fMRI data is to integrate multi-modality data into the analysis. It is therefore highly advantageous that TVDN is naturally able to handle multi-modality data. To understand this aspect intuitively, note that the stationary spatial features are by design modality invariant, and can be shared across multiple modalities. This imparts the TVDN framework with the ability to integrate information from both fMRI and MEG to estimate the spatial features. These shared spatial features will then be used to estimate the modality-specific temporal features, using information from both fMRI and MEG at each step, which will certainly improve estimation accuracy. While in this study we have shown how TVDN can operate seamlessly on both fMRI and MEG, we have not integrated the two for the current analysis because the data from paired samples are not available. Evaluating its performance on synchronized multimodality data would require larger collaborative studies involving both the fMRI and MEG centers.

One question of clinical interest is whether the dynamic RSFC predicts clinical outcomes, such as cognitive scores and disease risk. To address the question, the first and foremost step is to extract subject-specific dynamic RSFC features. However, the dynamic RSFC features from existing sliding-window methods give a set of RSNs of varying numbers across subjects, which makes it difficult to explicitly define unique spatial and temporal features for each subject. In contrast, TVDN extracts subject-specific dynamic RSFCs from both the fMRI and MEG data, which generates explicit spatial and temporal features that can be directly used to predict clinical outcomes. Evaluate the relationship between the dynamic RSFC features and clinical outcomes may potentially generate novel biomarkers for disease prediction.

### 3.1 Related methods

Sliding-window approaches are the most popular methods to extract dynamic RSFC from brain imaging data. However, these approaches do not typically allow for reconstructing the original brain signals in time or space, since they do not require a model of signal generation. And the temporal resolution of the inferred dynamic FC is inherently limited by the window length, which in turn is constrained by the requirement to have sufficient samples and signal-to-noise ratio within each window. In practice, this trade-off means that only slow changes in brain dynamics can be detected or tracked. Furthermore, in almost all current implementations, the sliding-window width is typically pre-specified and is not adaptable to the signal statistics or sampling noise in real time. In addition, they do not generate common features from the multiple modalities that may be available from a single subject (e.g. fMRI and MEG). This impedes information sharing across modalities and precludes benefiting from shared or redundant information between modalities.

Moreover, these methods typically suffer from very high data dimensionality, since at each window, the brain state is given by an entire network or several high-dimensional independent components - with no *a priori* notion of which features are actually evolving and which are static. Therefore the ability to detect discrete brain state switches then becomes dependent on the ability of unsupervised clustering algorithms like k-means or hierarchical clustering, to overcome the so-called”curse of dimensionality”. Finally, most dynamic extensions to static FC methods are purely data-driven and are not informed by biologically plausible modes of dynamicity in the brain, since they do not constrain which brain signal features can change dynamically and how - this aspect is discussed below. Therefore, sliding-window approaches present several limitations that must be overcome to gain further progress in critical neuroscience and clinical applications.

Several extensions of current methods have been proposed to address some of these limitations. To improve the sliding-window seed-based correlation approach, Faghiri et al. (2020) proposed a new metric replacing Pearson correlation between signals. Furthermore, Vergara et al. (2020) propose a robust method to determine the number of brain states from the sliding window methods. Hidden Markov models (Baum & Petrie, 1966) are an another robust alternative to capture brain state switches in the frequency domain. Vidaurre et al. (2017) and Quinn et al. (2018) used group level data to estimate the model parameters, assuming that study subjects share the same latent structure. Vidaurre et al. (2016) combined multivariate auto-regression model (Penny & Roberts, 2002) and hidden Markov model to obtain brain transitions, assuming that brain oscillations depend on the signals in a short time period prior to the current time. However, neither the improved sliding window methods nor the hidden Markov models are able to extract both the static spatial and dynamic temporal features. It is possible that further extension of these methods to account for both static and dynamic features will prove worthwhile, but out of scope of the current work.

### 3.2 Limitations and future directions

In our implementation, the tuning parameters of TVDN for fMRI were selected to minimize the average reconstruction error, and for MEG they were selected to minimize prediction error from cross validation among brain regions. A better tuning strategy might be to use cross-validation across individuals, where the data are split between training and testing individuals, and the tuning parameters are selected to minimize the prediction error in the testing individuals. However, because our switch detection relies on the entire time series of the whole brain, there is no existing method to split the study samples and validate parameter selection in the temporal switch detection procedure. Furthermore, a smaller prediction error may not necessarily imply a better prediction of disease states, like neurodegenerative disease risk. When the disease outcomes are available, an appropriate parameter selection strategy would be to select the tuning parameters that minimize the disease prediction error. Additional data and further research along these lines are ongoing in our laboratory.

Generally, fMRI signals have lower signal-to-noise ratio and temporal resolution than source-reconstructed MEG signals, which limits the former’s sample size available for parameter estimation. Hence, the brian state switching patterns extracted by TVDN from MEG appear clearer than from fMRI, with larger correlation with canonical networks. This suggests that MEG imaging could be a more informative technique to capture dynamic RSFC than fMRI. It is possible that alternative smoothing approaches than the one taken here might prove more effective on fMRI. It is possible that deconvolution of the hemodynamic response function might be helpful on fMRI data, an aspect that was not considered here.

## 4 Method

Dynamic RSFC contains separable spatial and temporal components (Geerligs et al., 2015). The spatial features of the dynamic RSFC capture the links among brain regions (Alexander-Bloch et al., 2010; Brier et al., 2014; Geerligs et al., 2015; Sanz-Arigita et al., 2010; Van Den Heuvel et al., 2009). The temporal features characterize the mental state evolvement of brain activities (Di et al., 2013; Gonzalez-Castillo et al., 2015; Kitzbichler et al., 2011; Moussa et al., 2011; Shirer et al., 2012). Furthermore, the spatial features are constrained by the stable brain structures, and hence they must be consistent over the signal sampling time and across the image modalities. Moreover, different modalities have distinct temporal resolutions, and therefore the temporal features are distinct across the modalities. Considering these characteristics of the spatial and temporal features, we develop a novel methodology to extract the time invariant spatial features and time varying temporal features.

### 4.1 Time-varying dynamic network

Let *X_i_*(*t)* be the brain signal at time *t* on the *i*th brain region of interest (ROI), 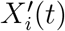 be the derivative *X_i_* with respect to *t* with *t* ∈ [0,*T*], representing the increment of brain activity at time *t*. Furthermore, let d be the number of ROIs, we write **X**(*t*) = {*X*_1_(*t*),…, *X_d_*(*t*)}^*T*^, and 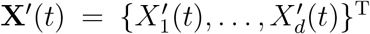. In practice, instead of the true signal, we observe a noisy signal at *n* discrete acquisition time points. Denote *t_j_* as the *j*th acquisition time, to accommodate the noisy data, we write

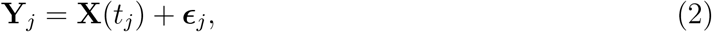

where **ϵ**_*j*_, *j* = 1,…, *n*, are independent mean zero random errors. Furthermore, we assume

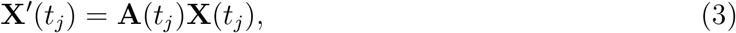

where **A**(*t_j_*) is a time-varying unknown matrix of size *d × d*. We name model (2) and (3) together the time-varying dynamic network (TVDN) model, where the dynamics of resting state functional connectivity is captured by the time-varying matrix **A**(*t*). Model (3) is a direct extension of the dynamic mode decomposition model (Brunton et al., 2016; Kutz et al., 2016). To see the connection, first note that (3) is equivalent to **X**(*t*_*j*+1_) = {**A**(*t_j_*)Δ*t* + **I**}**X**(*t_j_*), where Δ*t* is the unit measurement time and **I** is an identity matrix. When **A**(*t*) is a fixed matrix over time, we can treat **A**(*t_j_*)Δ*t* +**I** as a constant matrix. Then model (3) reduces to the dynamic mode decomposition model extensively studied in Brunton et al. (2016) and Kutz et al. (2016). Furthermore, when **A**(·) is a fixed matrix, model (3) is also a network diffusion model (Abdelnour et al., 2014), where 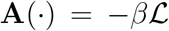 explains how brain activations from different ROIs are coupled together to generate new signals via the structural connectivity given by the matrix Laplacian 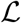 and the diffusivity constant *β*. Algebraic graph relationships have also been proposed, such that **A** may be given by the eigenvectors of structural (Abdelnour et al., 2018) or functional connectivity matrix (Becker et al., 2018), after a suitable transformation of the eigenvalues. While these approaches do not readily accommodate time-varying features of **A**, they point to an important property of the eigenvectors of **A**, which may be considered as resting state networks (RSN) (Abdelnour et al., 2018, 2014). Because these RSNs represent static brain connections or other nondynamic brain substrates, we propose the following constraints that together constitute the TVDN model:

1. The eigen-decomposition of **A**(*t_j_*) is in the form of

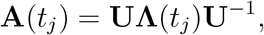

where we fix the eigenvector **U** but allow the eigenvalues to depend on time. Under this formulation, **U** may be considered as a set of spatial features that are stationary over time. The absolute magnitudes of the (time-varying) eigenvalues govern the relative importance of each of the RSNs.
2. We then impose the condition that dynamicity in RSFCs arise from discrete and potentially discontinuous shifts (“brain state switches”) in activity, which we accommodate by allowing the eigenvalues, i.e. diagonal elements of **Λ**(*t*), to be *piece-wise constant functions of time,* reflecting the phenomenon that the brain has a tendency to stay within, with sporadic cycling between the RSNs (Vidaurre et al., 2017). Therefore, let **τ**_0_ = (*τ_k_, k* = 0,1,…, *M, M* + 1; *τ*_0_ = 0, *τ*_*M*+1_ = *T*) be a set of true switching points, dividing the signal to *M* + 1 stationary segments, we write

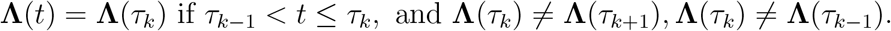 Such formulation suggests that when *t* ∈ (*τ*_0*k*-1_, *τ*_0*k*_], **Λ**(*t*) has a constant value at **Λ**(τ_0*k*_). And the values of **Λ**(·) are different at distinct time points that fall into two consecutive segments constructed by the switching points.
3. The number of nonzero eigenvalues in **Λ**(·) represents the number of intrinsic mental states in the task free brain activity data. It is well known that only a few RSNs are typically operational in the brain, and canonical RSFC can be well captured by 7-20 such RSNs (Yeo et al., 2011). In prior graph theoretic models also, **A**(·) are assumed to be low rank matrices (Abdelnour et al., 2018; Raj et al., 2019). It is therefore plausible to assert that the number of such RSNs or brain sates is quite small. Hence our final constraint is that

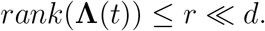

### 4.2 Model interpretations

Since **A**(*t*) is constant in a given segment, the solution of the ordinary differential equation (3) in the k-th segment is given by **X**(*t*) = **U**exp{**Λ**(*t*)}**U**^-1^ **X**_0*k*_, where **X**_0*k*_ is the initial value at the kth segment and **Λ**(*t*) is a constant matrix in the kth segment. Let us define the real and imaginary components of the *j*-th eigenvalue as *λ_j_* = *γ_j_* + *i*2*πf_j_*. Then the underlying signal in the *k* segment satisfies

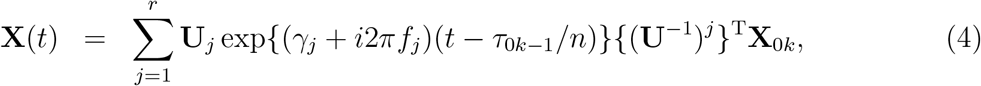

where (**U**^-1^)^*j*^ is the jth row of **U**^-1^. The real term *γ_j_* is interpreted as a coefficient that determines the growth or decay of the signal during this segment, and the imaginary component *f_j_* is interpreted as the oscillation frequency of the mode (Kunert-Graf et al., 2019) in cycles per sample interval, which is 2 seconds for fMRI data and 1/60 seconds for MEG data). Therefore, when an estimator for **Λ**(*t*), say 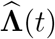, is available for the kth segment, we can directly infer the grow/decay constant in the segment as 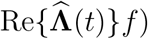 and the signal frequency as 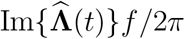.

It is worth mentioning that in situations where mulitple modalities are available for the same subject, e.g. fMRI and MEG, the spatial features **U** may be considerd to be shared between the modalities. In those cases TVDN will be able to aggregate the signals to generate a common estimator for **U** across the modalities. Then this common estimator will be used to obtain the modality specific temporal features **Λ**(*t*), where the information borrowing is clearly embedded in the estimation through sharing **U**. Augmentation with multi-modal data can potentially improve estimation accuracy.

### 4.3 Estimation of the spatial and temporal features

We estimate the spatial feature **U** through a kernel based method and detect the critical points for brain state switches via a switch detection algorithms.

#### 4.3.1 Notation

Let *t*_1_,…, *t_n_* denote *n* signal acquisition time points. We denote **M**_*r*_ and **M**_*r*×*r*_ be the first *r* column and the first *r × r* block of **M**, respectively. Furthermore, define 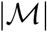 be the cardinality of an arbitrary set 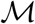. Let ||**M**||_*F*_ be the Frobenius norm of matrix **M**. For a vector a, let ||a|_1_, ||a||_2_ be its *L*_1_,*L*_2_ norm, respectively. Let **M**^−1^ be the generalized inverse of matrix **M**.

#### 4.3.2 B-spline smoothing

To obtain a proper estimator for *X*(*t*) and *X*’(*t*) from the noisy observation *Y*(*t*), we first de-noise the signals through the B-spline smoothing as follows

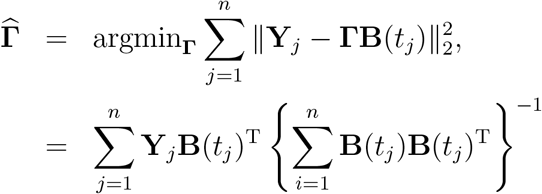

and

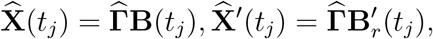

where **B**(·) is the *b*th order B-spline basis with *N* interior knots and 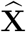 and 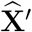 are the smoothed version of **X** and **X**’, respectively.

#### 4.3.3 Estimation of the spatial features U

On these smoothed signals, we estimate at any time point of interest *t_s_* the matrix **A**(*t_s_*) by minimizing

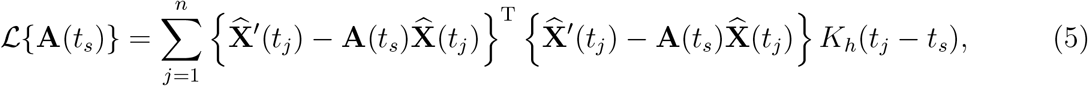

where *K_h_*(|*x*|) = 1/*hK*(|*x*|/*h*) is a kernel function with *h* be the bandwidth. Here *K_h_*(|*x*|) is a deceasing function of |*x*|. Hence when estimating **A**(*t_s_*), *K_h_*(|*x*|) weighs the samples higher when they are closer to *t_s_*. The width h of the kernel controls how “local” the estimator of **A**(*t*) is; if it is large, then the estimator of **A**(*t*) would hardly change over time, and it would reduce to the DMD model (Kunert-Graf et al., 2019). A typical choice of *K_h_*(|*x*|) is the Gaussian density function with standard deviation *h*. The bandwidth h is often selected to satisfy *h* → 0 when *n* → ∞ so that even if n grows, when estimate **A**(*t_s_*), the amount of information used in the estimation remains fixed. In Figure 8, we show the weights *K_h_*(*t* — 180) across *t* ∈ [1, 360] in seconds for fMRI and weights *K_h_*(*t* — 30) across *t* ∈ [1,60] in seconds for MEG when the rule-of-thumb bandwidth h (page 48 in Silverman (1986)) was selected.

**Figure 8:**
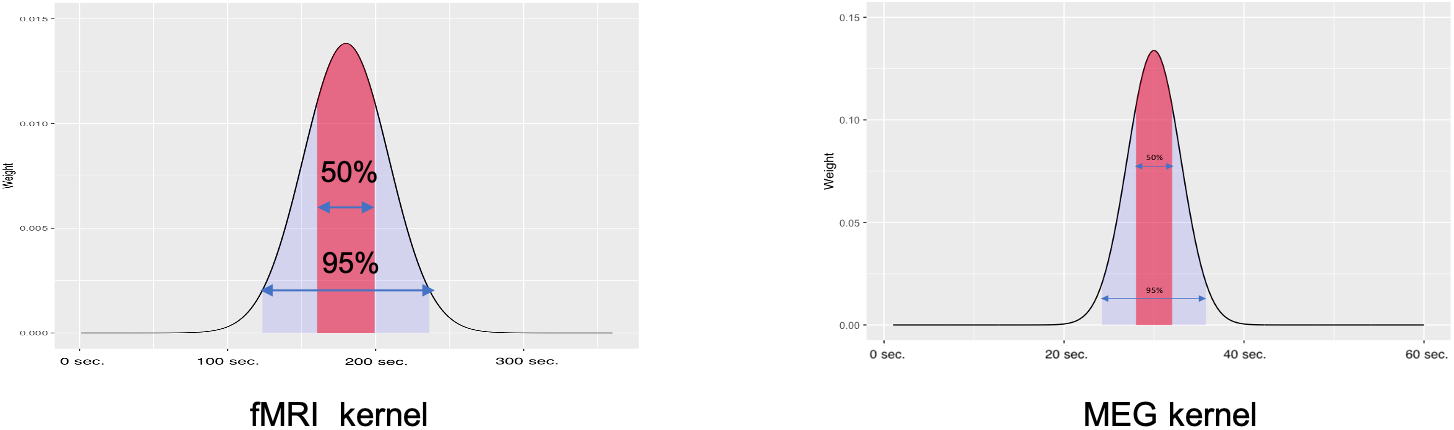
The weights *K_h_*(*t* — 180) across *t* ∈ [1,360] in seconds for fMRI (left) and weights *K_h_*(*t* — 30) across *t* ∈ [1,60] in seconds for MEG (right) when the rule-of-thumb bandwidth *h*.

Minimizing 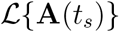 has a close form solution for **A**(t_s_) as

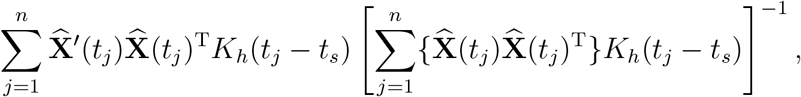

which is the Nadaraya-Watson estimator (Nadaraya, 1964; Watson, 1964) regularly used to estimate functions at specific time points.

To account for the fact that 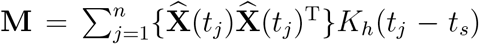 can be a low rank matrix, we replace the matrix inverse above with a truncated-rank inverse such that all eigenvalues of **M** below a threshold value are set to zero and removed from the pseudoinverse. Formally, define a truncation function *ρ_λ_* as 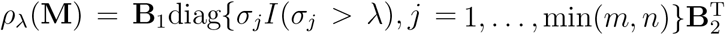, where **B**_1_, **B**_2_ are the left and right singular vectors and *σ_j_* is the *j*th singular value of **M**. Herex we choose a truncation threshold *b*_0_ = *O*(*h*^2^*r* + *n*^-1/2^*N*^-1/2^*h*^2^*d*), which is in the order of 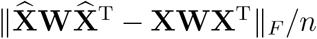 with **W** = diag{*K_h_*(*t_j_* — *t_s_*), *j* = 1,…, *n*} for a specific time point *t_s_*. Here, the first order *h*^2^*r* comes from the kernel smoothing and the second term *n*^-1/2^*N*^1/2^*h*^2^*d* comes from the B-spline smoothing. When *n* → ∞ and *n*^-1/2^*N*^1/2^ → 0 as *n* → ∞, the error 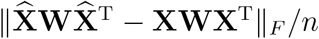 and in turn the quantity *b*_0_ go to 0 when sufficient samples are collected overtime.

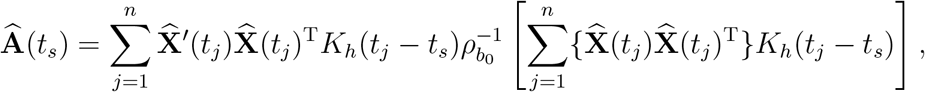

The truncation function is specially designed for the high correlated sequences where the **XWX**^T^ is a low rank matrix, but 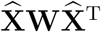 can be full ranked due to the additional estimation error 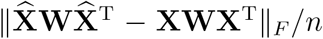. Using the truncation function helps to remove spurious eigenvalues, which not only improves the estimation accuracy but also stabilizes the computation.

When **X** is full row rank matrix, we can show that 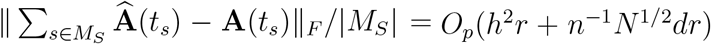, which goes to 0 when sample size increases under mild conditions. Here, the first term *h*^2^*r* is the order of the estimation error in the kernel regression procedure and the second term *n*^-1^*N*^1/2^*dr* is the order of the error from the B-spline smoothing procedure. When *h* → 0 and *n*^-1^*N*^1/2^ → 0 as *n* → ∞, the estimation error of 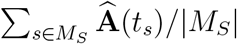 vanishes along with the increment of the sample size. This suggests that we need sufficient samples to recover the underlying true parameters. This fact also explains the phenomenon in the real data analysis that the reconstruction of the MEG signals is better than that of the fMRI signals. Therefore, we extract the estimator for **U** as the eigenvector of 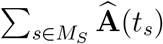, denoted as 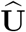.

#### 4.3.4 Brain-state switch detection

Define **M**_*r*_. as the first r rows of matrix **M**, and **M**_*r×r*_ as the first *r × r* block matrix of **M**. Because

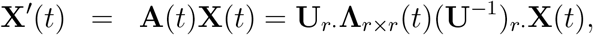

after obtaining 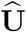 we reduce the data dimension as

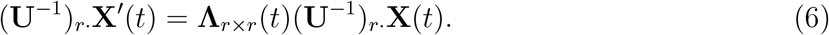

It is worth mentioning that multiplying both sides by (**U**^-1^)_*r*_.**X**(*t*) projects the *d* dimensional ROIs to a lower *r* dimensional space. Furthermore, **Λ**_*r×r*_(*t*) is a diagonal matrix, which contains only *r* unknown parameters. Such dimension reduction procedure is crucial for speeding up and stabilizing the switch detection algorithm, which makes the brain state switch detection practically feasible for signals from a large number of ROIs.

Let 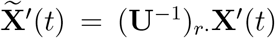 and 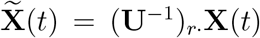, we obtain the estimator for true switch number *M* and locations *τ_k_*’s through minimizing a modified Bayesian information criteria defined as

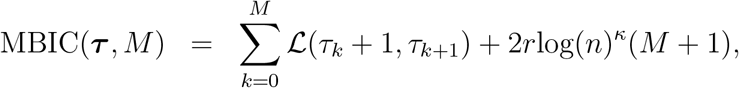

where *κ* is a constant, and

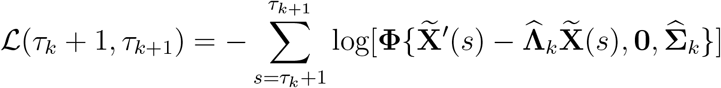

with **Φ**(**x, a, s**) be the multivariate normal density function with mean **a** and variance covariance matrix **s** evaluated at **x**. To solve the minimization problem, we iterate over all possible segmentations of the sequence. For the samples in a given segment, say *s* ∈ (*τ_k_* + 1, *τ*_*k*+1_], we obtain 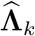 as

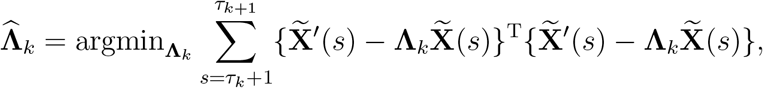

subject to the fact that **Λ**_*k*_ is a *r × r* diagonal matrix, and obtain 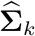 as the estimated sample covariates defined as

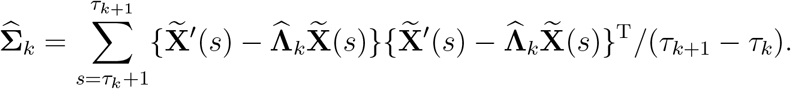

We then find the best segmentation, that is the best (*τ, M*) that minimizes *MBIC*(*τ, M*) as the estimated locations and number of the switch points. In short, we obtain the estimators as

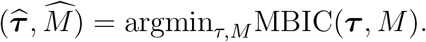

We employ the dynamic programming algorithm as detailed in Dynamic programming algorithm as follows (Jackson et al., 2005; Killick et al., 2012) to efficiently evaluate all possible segmentations and obtain 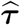 and 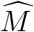. The dynamic programming algorithm finds the optimal value recursively, avoiding re-computing the 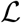 over overlapped segments (Bellman & Roth, 1969; Bement & Waterman, 1977; Du et al., 2016a; Yau & Zhao, 2016). The computational cost is *O*(*n*^2^*M*_*max*_) and storage is *O*(*nM_max_*), where *M_max_* is the maximum number of switches points in the signal (Du et al., 2016b).

#### 4.3.5 Tuning parameter selection

We choose rank r so that the first *r* modulus of the eigenvalues of 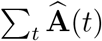 comprise 80% of the total sum of them, where the summation is taken over a random subset of times. Furthermore, we select the bandwidth for kernel in (5) to be the rule-of-thumb bandwidth times 0.5. Moreover, for the resting state fMRI data, we select *κ* to minimize the variation of the number of switches across the subjects. For the resting state MEG data we select *κ* through resampling over acquisition time. More specifically, for a given *κ*, we select five subsamples, where the *j*th sample contains data at times 5*t* + *j* with *j* = 1,…, 5 and *t* = 1,…, (*n* — 5)/5. Since the five sequences are in conjunction with each other, they should have similar numbers of switches.

##### Dynamic programming algorithm

Input: (1) *L_min_*, the minimum distance between two change points; (2) *M_max_*, the maximum number of change points; (3) *κ* value.

1. For 0 ≤ *i* ≤ *n* — *L_min_* + 1 and *i* + *L_min_* ≤ *j ≤ n* + 1, calculate 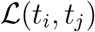.
2. Initialize 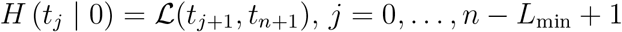.
3. For 1 ≤ *s* ≤ *M_max_* and 0 ≤ *i* ≤ *n* — *s* — *L_mon_*, update

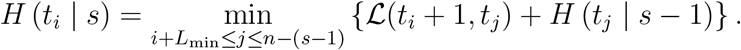 Record the locations of *s* change points that yield *H* (*t*_0_ | *s*), denoted by 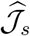.
4. For 1 ≤ *s* ≤ *M_max_*, find

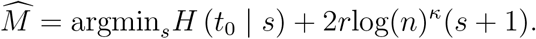 The corresponding estimated switch point set is 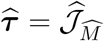.

Output: 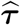 and 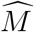.

### 4.4 Sliding window approaches

We implement TVCOR, TVPCA and TVDMD as follows where we construct windows in different sizes sliding by 4 frames in each step. Let 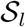 be the set of time point indices in the sliding-window *l, l* = 1,…, *L*. For matrix 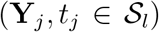, TVCOR calculates the pairwise correlation between signals from different ROIs, TVPCA extracts the principle components, TVDMD extracts the dynamic modes (Brunton et al., 2016; Kunert-Graf et al., 2019) from the brain signals. Next we vectorize the resulting correlations, principle components and dynamic modes and cluster them into four clusters, corresponding to the number of true segments in the simulation. Finally, we obtain the switch locations as the time points where the vectorized correlations, principle components and dynamic modes switch the cluster memberships.

### 4.5 Simulation method

We construct **A**(*t*) = **UΛ**(*t*)**U**^-1^ for *t* = 1,…, 180, where **Λ**(*s*) are singular vectors and **U** are eigenvalues and eigenvectors estimated from a functional magnetic resonance imaging (fMRI). Here **Λ**(*s*) is a rank six matrix, which contains three switches at the 50, 99,144th time points. We simulate data from model (2) and (3), where *ϵ_j_* = **UΛ**(1)**U**^-1^*ξ_j_*10, *j* = 1,…, *n* and *j* is a sparse error vectors with 10% nonzero entries. Each nonzero element in *ξ*(*t*) is independently generated from a normal distribution with standard error (*t*_2_ — *t*_1_)/8. We simulate the data 100 times with the same set of parameters. Then we implement TVDN to obtain the estimated spatial feature 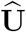, the switch locations and the temporal features 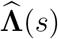, where we select *κ* = 1.53 throughout the simulations to achieve satisfactory results.

In the left panel of Figure 9 (a), we plot the reconstruction errors defined as

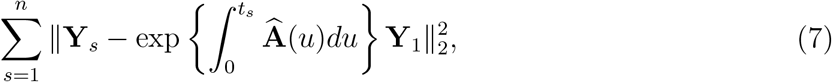

when different ranks for 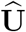 are selected in the estimations. The results show that the estimation error substantially drops from the setting when *r* = 4 to the setting when *r* = 6. Furthermore, when *r* > 6, the reconstruction error starts to incline. The convex phenomenon attributes to the tradeoff between the dimension reduction described in (6) and switch detection accuracy: when selecting a larger r, the transformation (**U**^-1^)_*r*_.**X**(*t*) contains more information in **X**(*t*), but it increases the estimation errors for the brain state switch detection algorithm; on the other hand, selecting a smaller r improves the switch detection accuracy, but (**U**^-1^)_*r*_.**X**(*t*) contains less information in the original data. On the right panel in Figure 9 (a), we show the MBIC values when selecting *κ* = 1.53, which reaches the minimum when three switches are selected.

We select six (the rank of **A**(*s*)) principle components and dynamic modes throughout the simulations. Figure 9 (b) plots the distribution of the Hausdorrff distance between the true and estimated switches for the different methods. A smaller Hausdorrff distance implies a better estimation. It can be seen that the switches from TVDN have the smallest Hausdorrff distance with the truth. There are several occasions that the sliding-window approaches outperform the TVDN method. This is because we specify the true number of segments in the sliding-window approaches, while we leave this parameter unknown in the TVDN approach and allow TVDN to choose it adaptively. To illustrate the pattern in more details, we plot the resulting switch locations from the sliding-window methods in Figure 9 (c). Figure 9 (c) shows that none of the three methods correctly identify the switches. In addition, the sliding-window based methods are sensitive to the window size changes, which leads to substantial different results when varying the window sizes.

**Figure 9:**
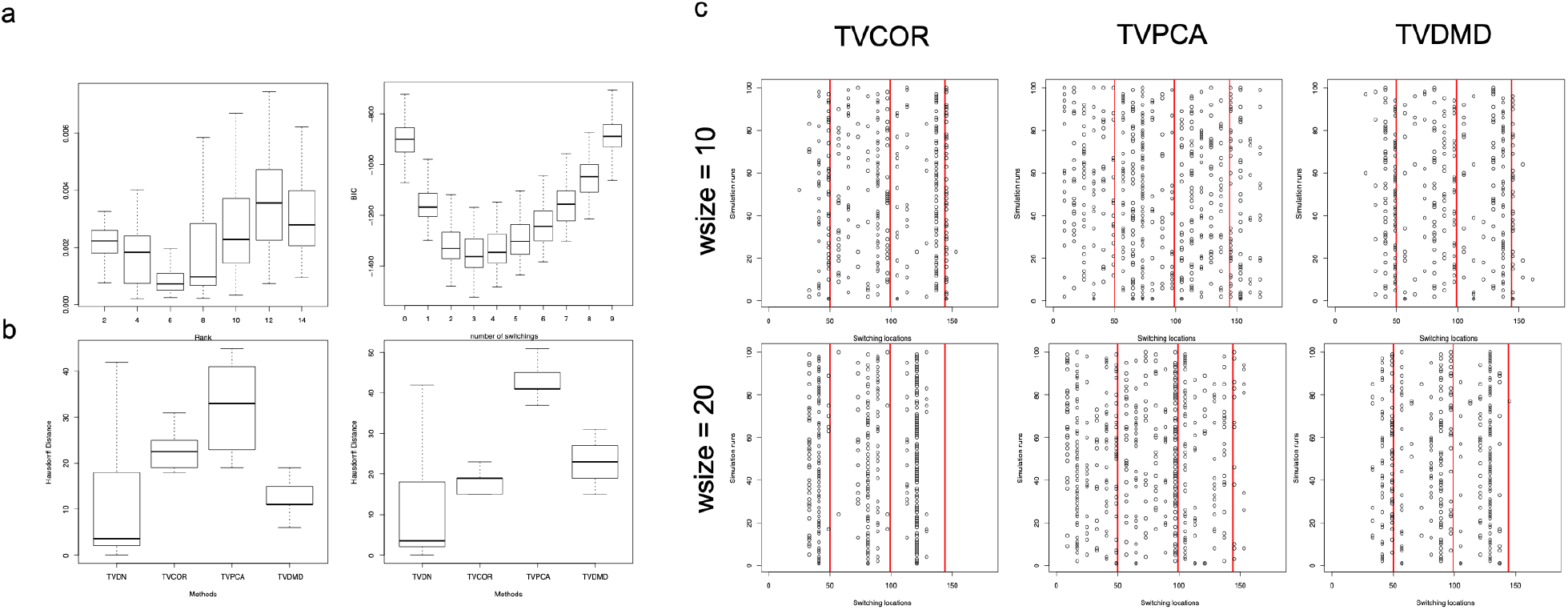
The simulation results with three switches. Here **A**(*t*) has six ranks. (a) Left: the boxplot of the reconstruction error in (7) from 100 simulations when choosing different ranks for 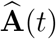 in the estimation. Right: the boxplot of the MBIC values at different number of switches when *κ* =1.53. (b) Hausdorrff distance between true switches and the estimated switches. The window sizes (wsize) selected are 10 (left) and 20 (right) for the sliding-window based methods. (d) Chang point locations for window sizes 10 (top) and 20 (bottom) for the TVCOR (left), TVPCA (middle) and TVDMD (right) methods.

Finally, we adopt the same simulation procedure while assume a time invariant **A**(*s*). Then we implement TVDN on the resulting high dimensional sequences and reconstruct the observed data. We show the mean of the estimators and 95% confidence intervals in Figure 10 in Appendix. The results show that even if **A**(*s*) is stationary over time, TVDN correctly extracts the spatial and temporal features from the brain signals.

### 4.6 Data and preprocessing

#### 4.6.1 fMRI data

Resting state fMRI data from 103 health subject was acquired at the UCSF Neuroimaging Center using a Siemens 3T TIM TRIO scanner using a T2*-weighted AC-PC aligned echo planar imaging (EPI) sequence with the following parameters: TR = 2000 ms, TE = 29 ms, flip angle = 75, FOV = 240 x 240, slice thickness = 3.5 mm. Each fMRI was recorded over six minutes with 0.5 HZ sampling rate. Preprocessing included slice-timing correction (Cox & Hyde, 1997), image realignment to correct for motion (Jenkinson & Smith, 2001), and intensity normalization. The head-motion parameters were estimated before any spatiotemporal filtering was used (Jenkinson et al., 2002). After regression of nuisance signals, fMRI was coregistered on the T1-weighted anatomical image, and the resulting time-series were normalized to MNI space with the non-linear registration from ANTS (Avants et al., 2009).

Following time series extraction, data were detrended and a bandpass filter was applied between 0.009 and 0.08 HZ. To remove the boundary effect from the filtering procedure, we remove the first 25 sampling points. Hence, the total length of the signal is 155.

#### 4.6.2 MEG data

MEG data was acquired in the Biomagnetic Imaging Laboratory at University of California, San Francisco (UCSF) with an Omega 2000 whole-head MEG system from CTF Inc. (Coquitlam, BC, Canada) with 1200 Hz sampling rate. For resting state data analysis, two subjects were instructed simply to keep their eyes closed and stay awake. We collected 4 trials per subject, each trial of 1-min length with a sampling rate of 1200 Hz. We randomly chose 10 seconds or equivalently 12000 time samples for brain source reconstructions for each subject. Additionally, for one subject, MEG data was collected across two sessions for a an eye-opening-closing task. To measure eye opening and closing, two pairs of Electrooculography (EOG) electrodes were placed to the left and right of the eye during MEG scans. A potential difference was recorded when the subject blinked eyes and a signal peak occurred in the EOG channel of scanned data. We manually labeled EOG peaks to indicate time periods of eye opening and closing for TVDN analyses. Across both resting and eye-opening task, all MEG sensor locations were co-registered to each subject’s anatomical MRI scans. The leadfield for each subject was calculated in NUTMEG (Dalal et al., 2004) using a single-sphere head model (two spherical orientation leadfields) and an 8 mm voxel grid. Each column was normalized to have a norm of unity. The data were digitally filtered to remove DC offset and any other noisy artifact outside of the 1 to 45 Hz bandpass range.

To infer the neuronal activity in the source space from the MEG recordings, which were in sensor space, source localization was performed using time-frequency optimized adaptive beamforming (Dalal et al., 2004) using the custom-built open source NUTMEG software tool. Since this study focuses on the cortical areas and only the sources belonging to the 68 cortical regions were selected based on the Desikan-Killiany parcellations. The time-course of activity in each of the 68 brain regions was estimated by averaging the time-course of source activity estimated from voxels within a 20 mm radius of its centroid.

The resting state MEG data was down sampled to 600 HZ, while the eye-opening-closing MEG was down sample to 1200 HZ in our analysis.

## Appendix

## A Additional simulation results

**Figure 10:**
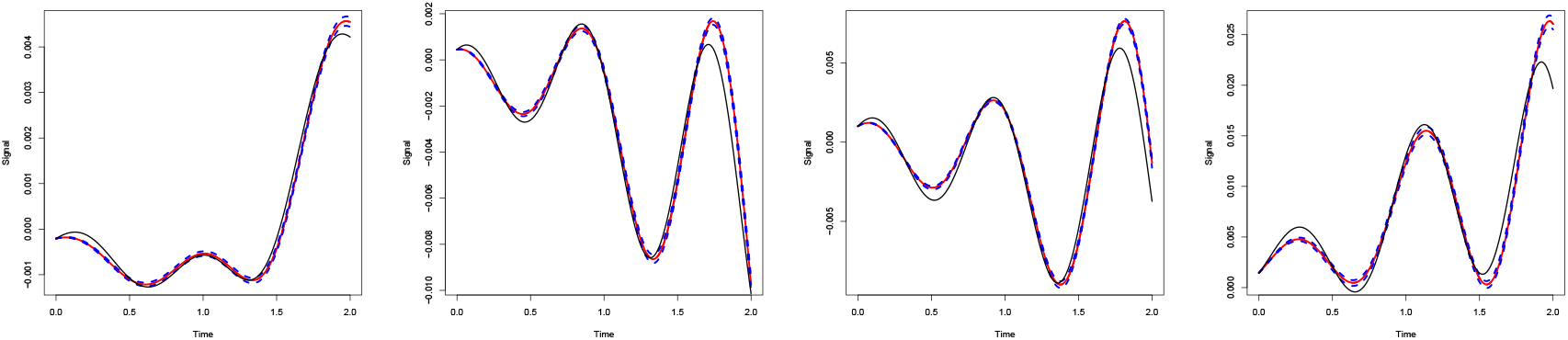
The simulation results with no switch. Here **A**(*t*) has six ranks. The black lines are the true **X**(*t*) at four selected regions. The red solid and dash curves are the mean and median of the estimators and above and below blue curves are the 95% empirical confidence intervals. The figures from left to right represent the results of the estimators whose mean squared errors follow on the 0%, 25%, 50%,75% quantiles of the mean squared errors across all simulations. The confidence intervals are narrow because the simulated random errors have small variabilities.

## B fMRI additional results

**Figure 11:**
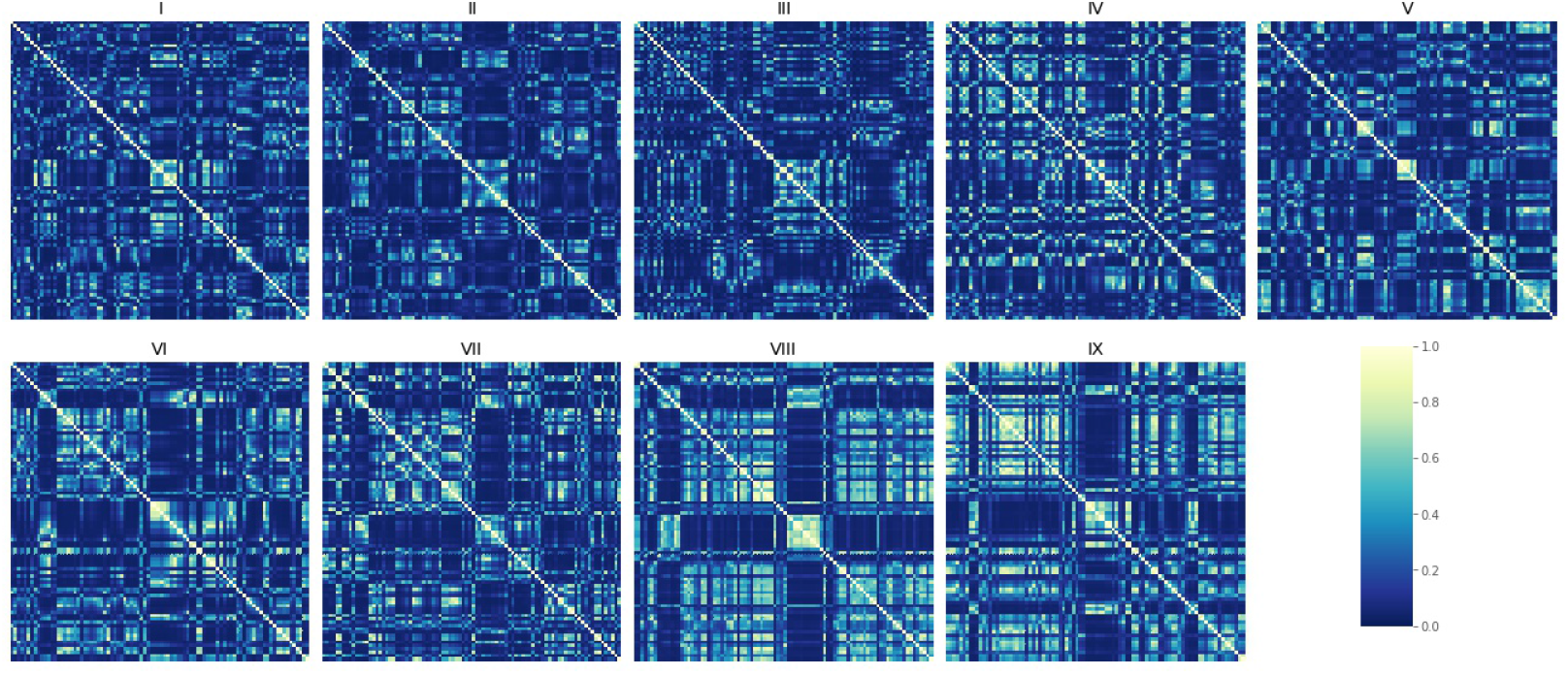
The pairwise connectivity over 90 ROIs from the first fMRI example.

**Figure 12:**
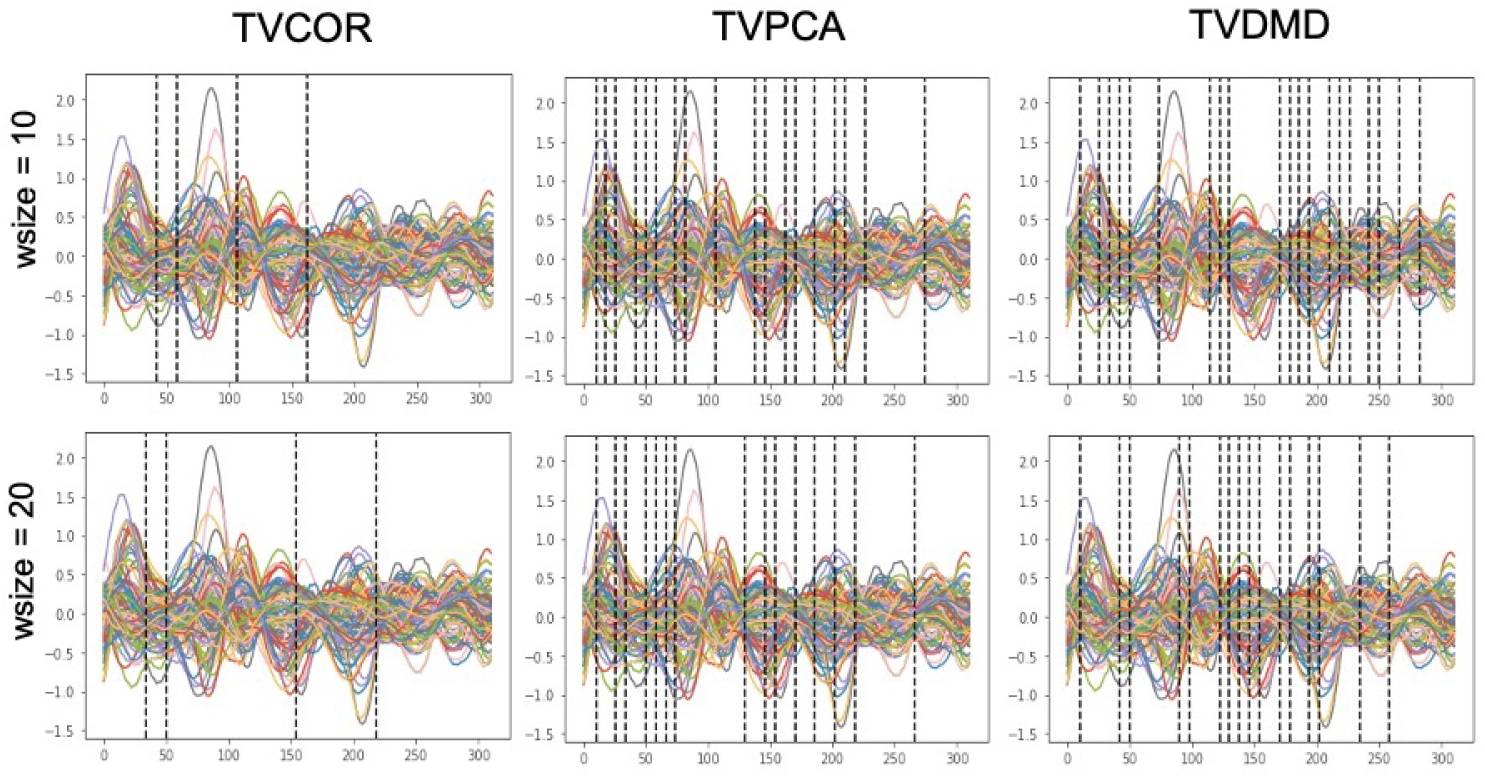
The brain state switch detection is not robust across different window size selections for the sliding-window approaches on the first fMRI example.

## C MEG additional results

**Figure 13:**
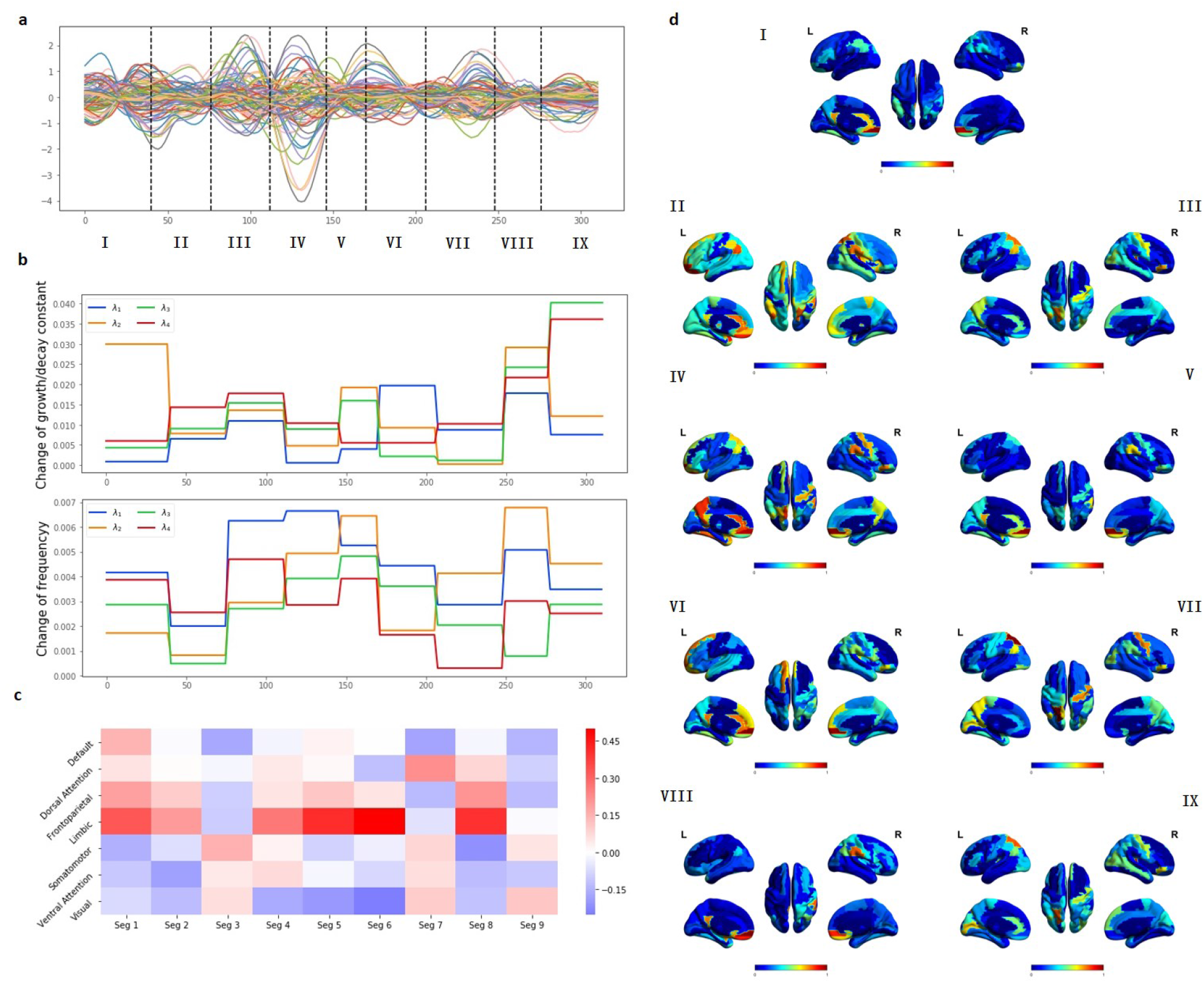
The results from the second fMRI sample. (a) The the real sequences with switch locations detected by TVDN. The horizontal line is the acquisition time. (b) Changes of growth/decay constant 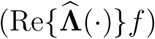, changes of the frequencies 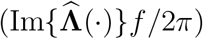. (c) The Pearson correlation between the weighted spatial features and the seven canonical networks. (d) The weighted spatial features across different segments detected by TVDN.

**Figure 14:**
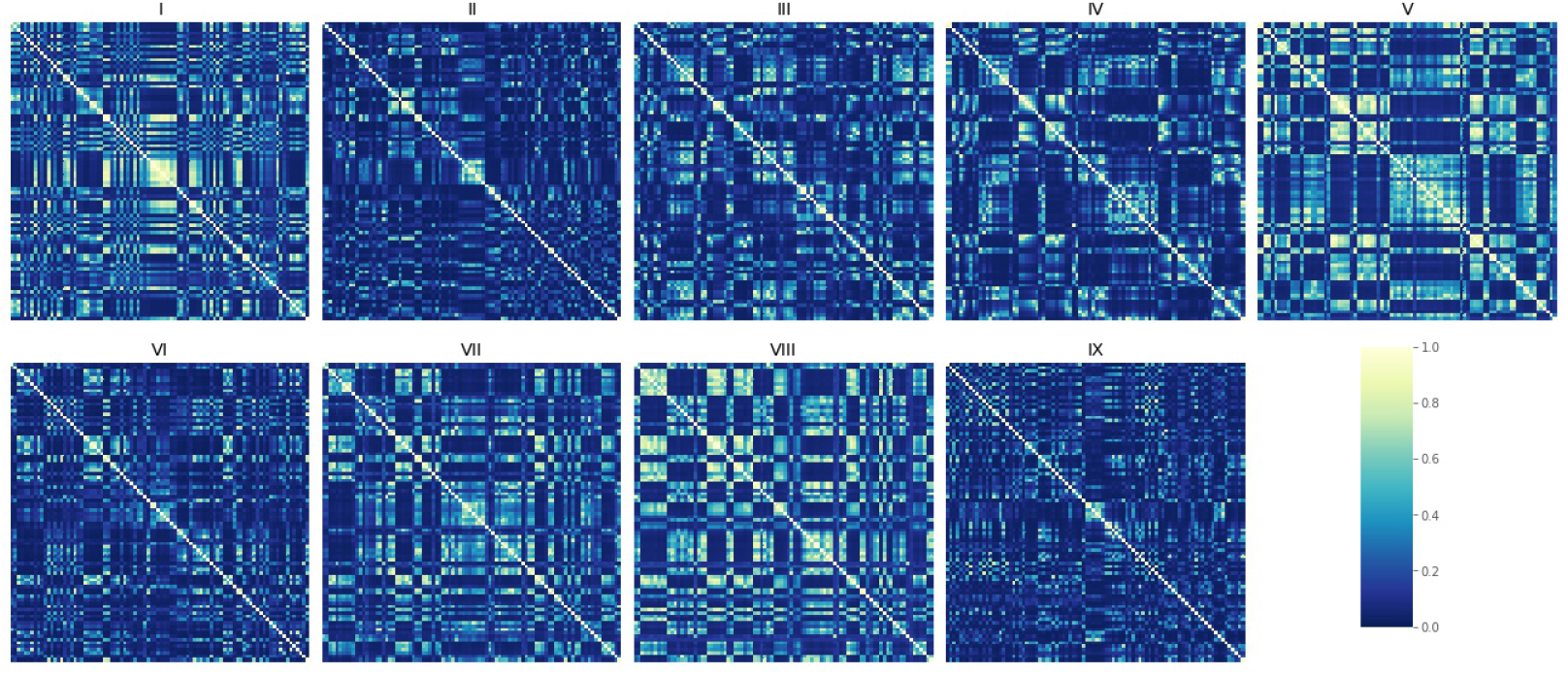
The pairwise connectivity over 90 ROIs from the second fMRI example.

**Figure 15:**
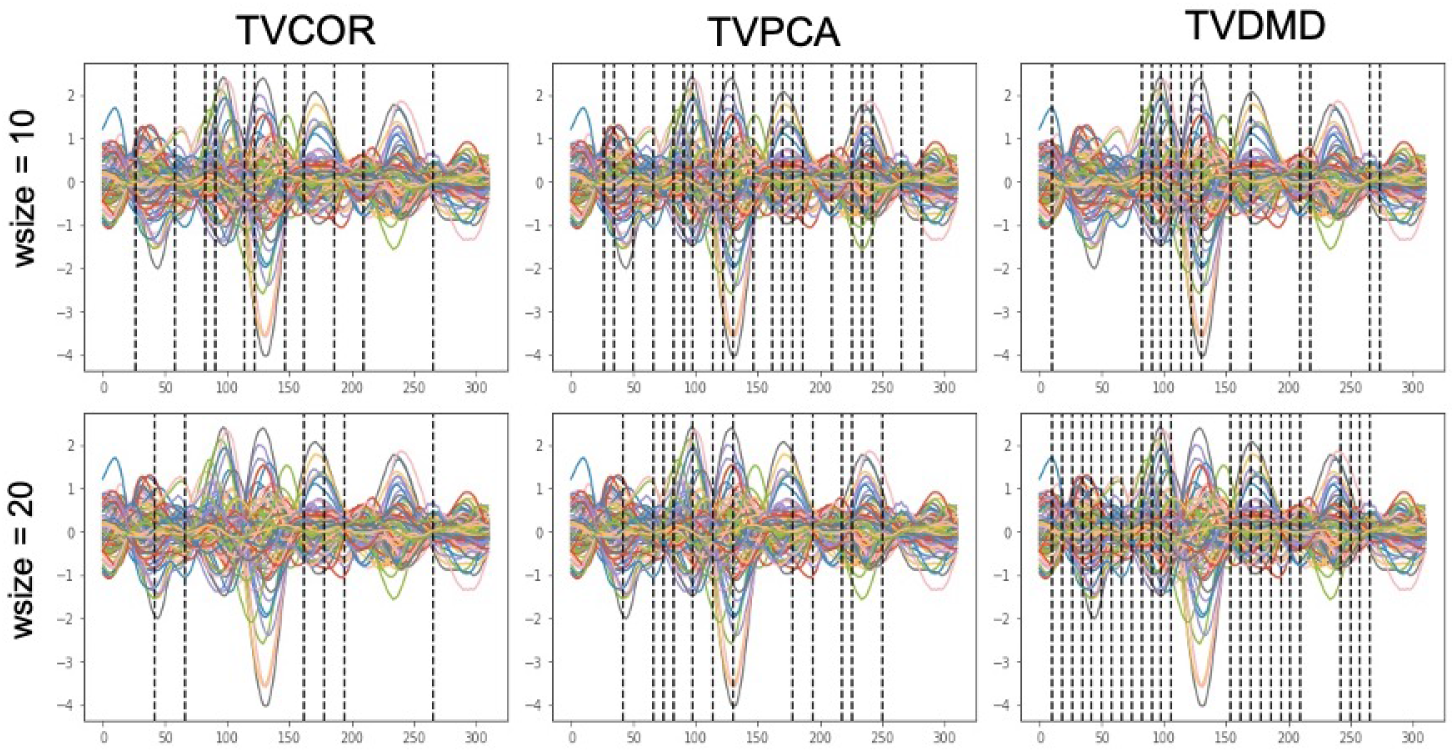
The brain state switch detection is not robust across different window size selections for the sliding-window approaches on the second fMRI example.

**Figure 16:**
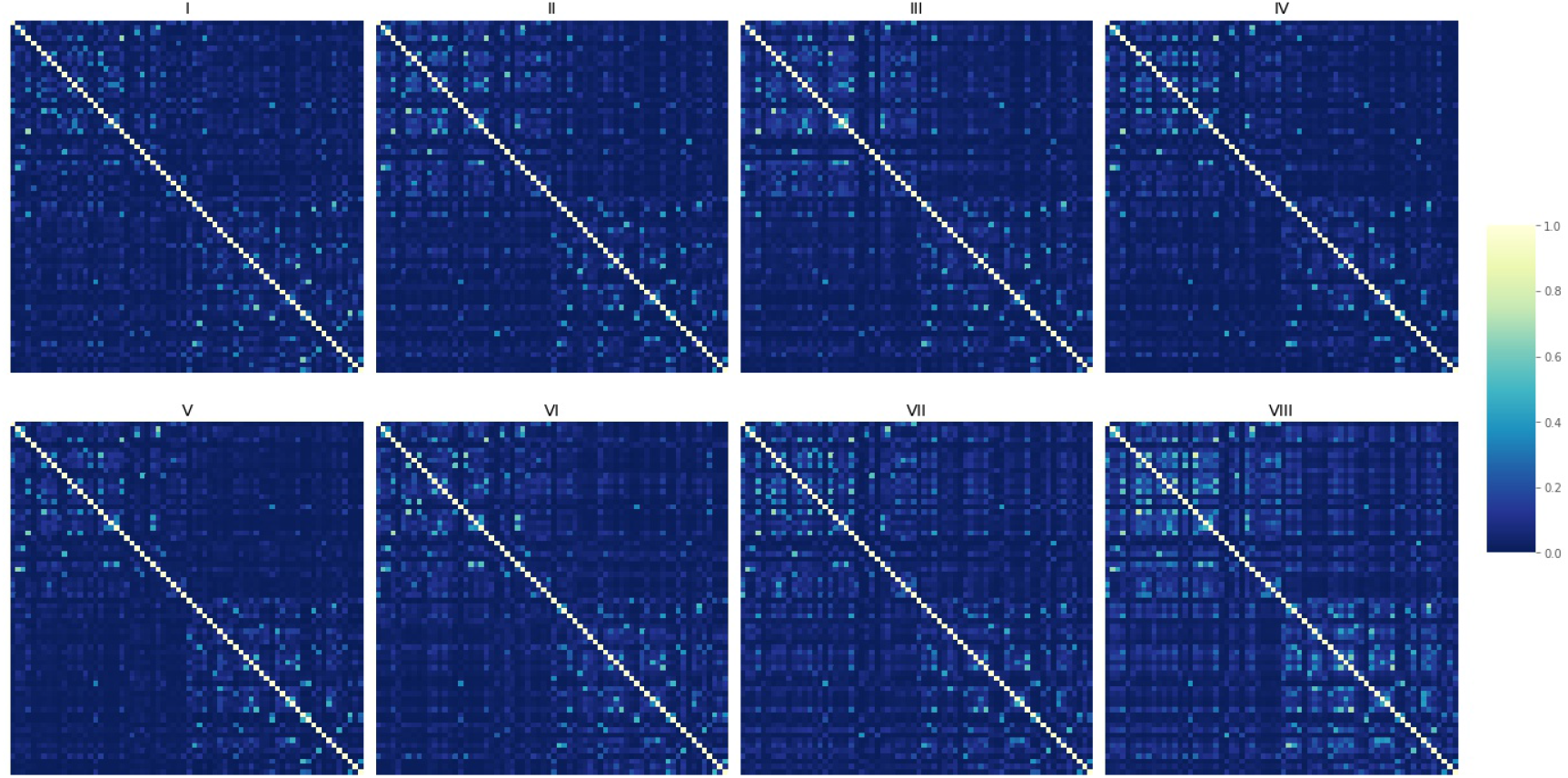
The pairwise connectivity over 68 ROIs from the first MEG resting state example.

**Figure 17:**
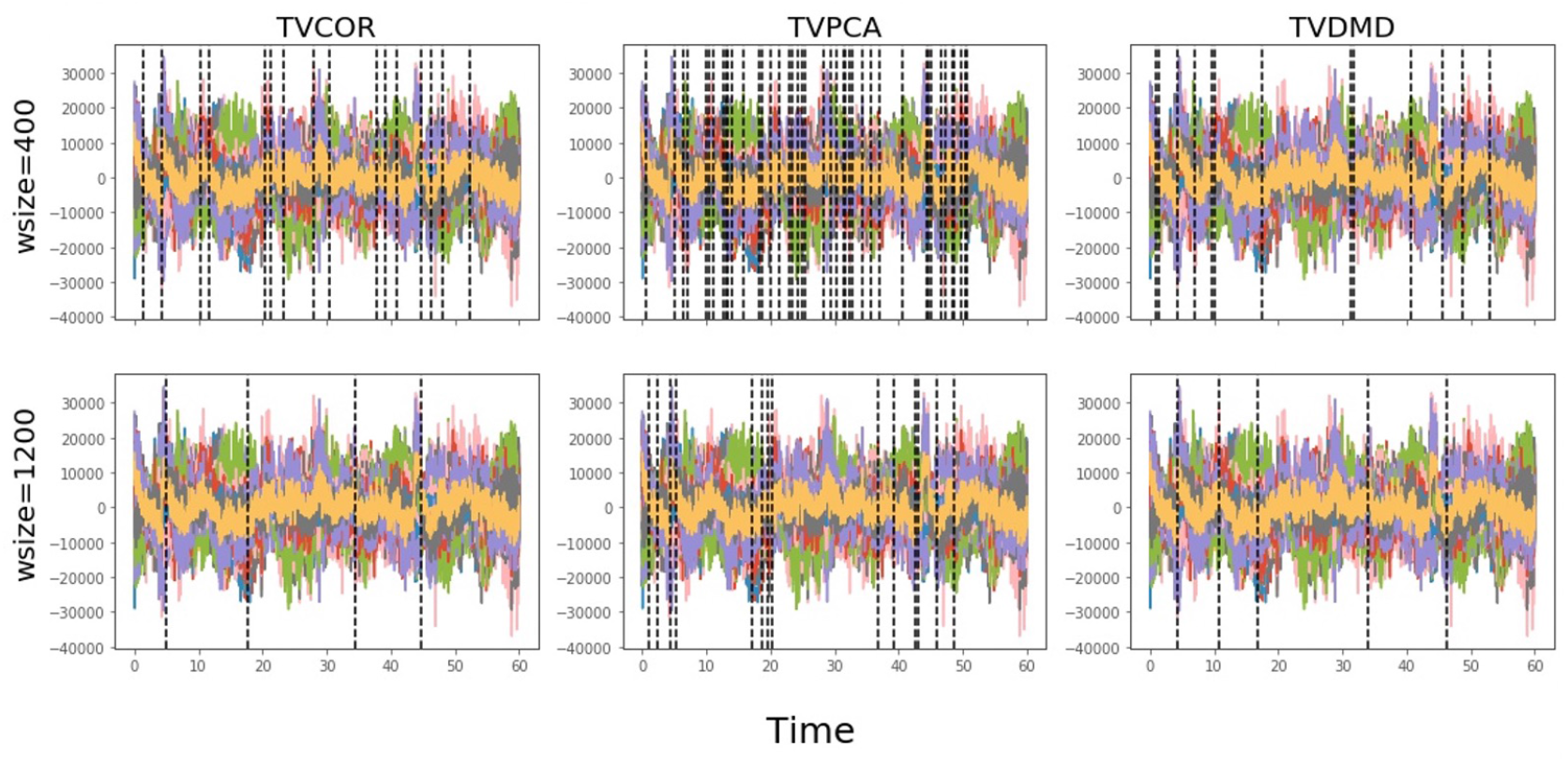
The brain state switch detection is not robust across different window size selections for the sliding-window approaches on the first MEG resting state example.

**Figure 18:**
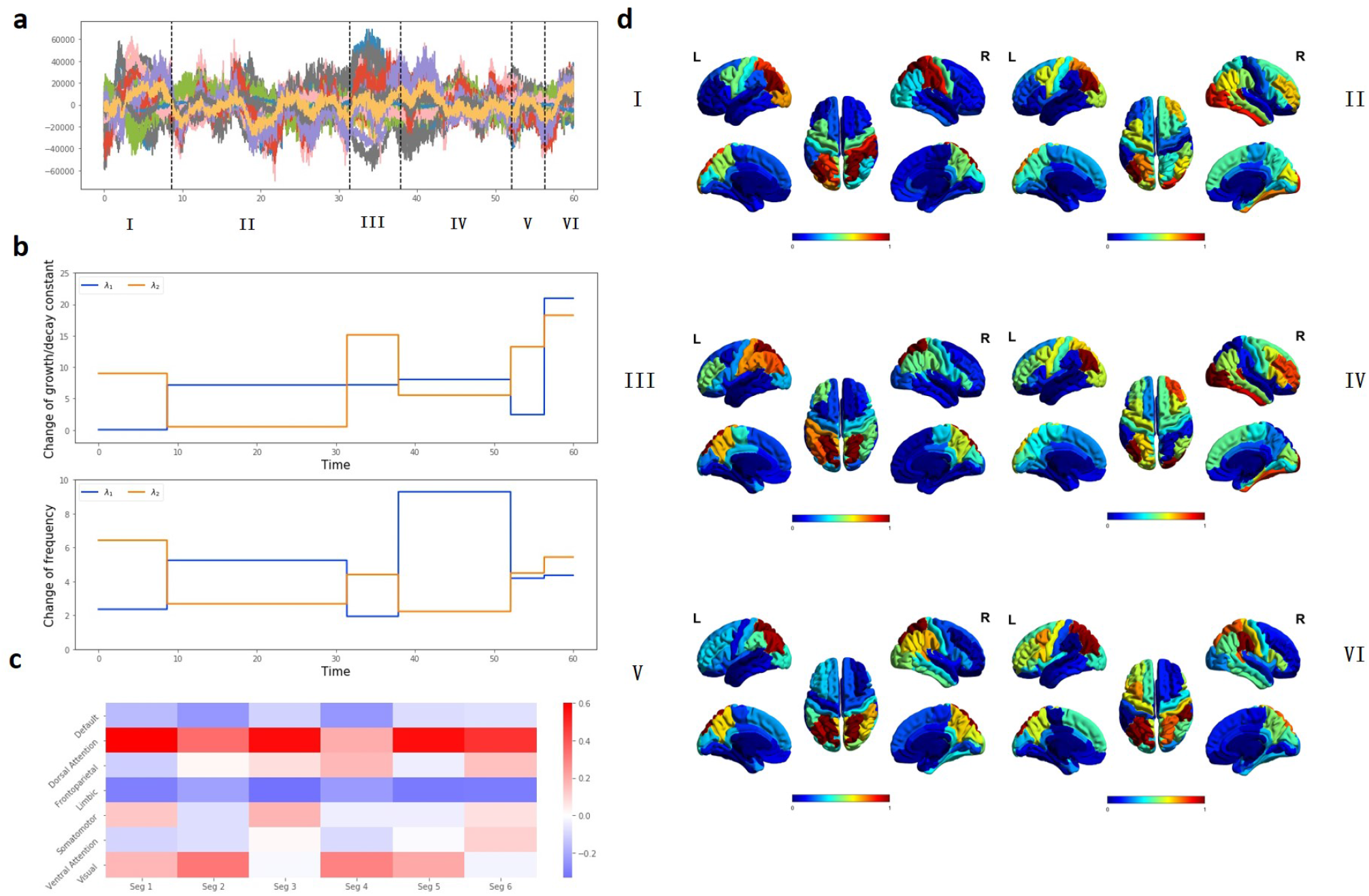
The results from the second resting state MEG record. (a) The the real sequences with switch locations detected by TVDN. The horizontal line is the acquisition time. (b) Changes of growth/decay constant 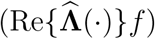, changes of the frequencies 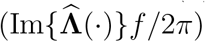. (c) The Pearson correlation between the weighted spatial features and the seven canonical network. (d) The weighted spatial features across different segments detected by TVDN.

**Figure 19:**
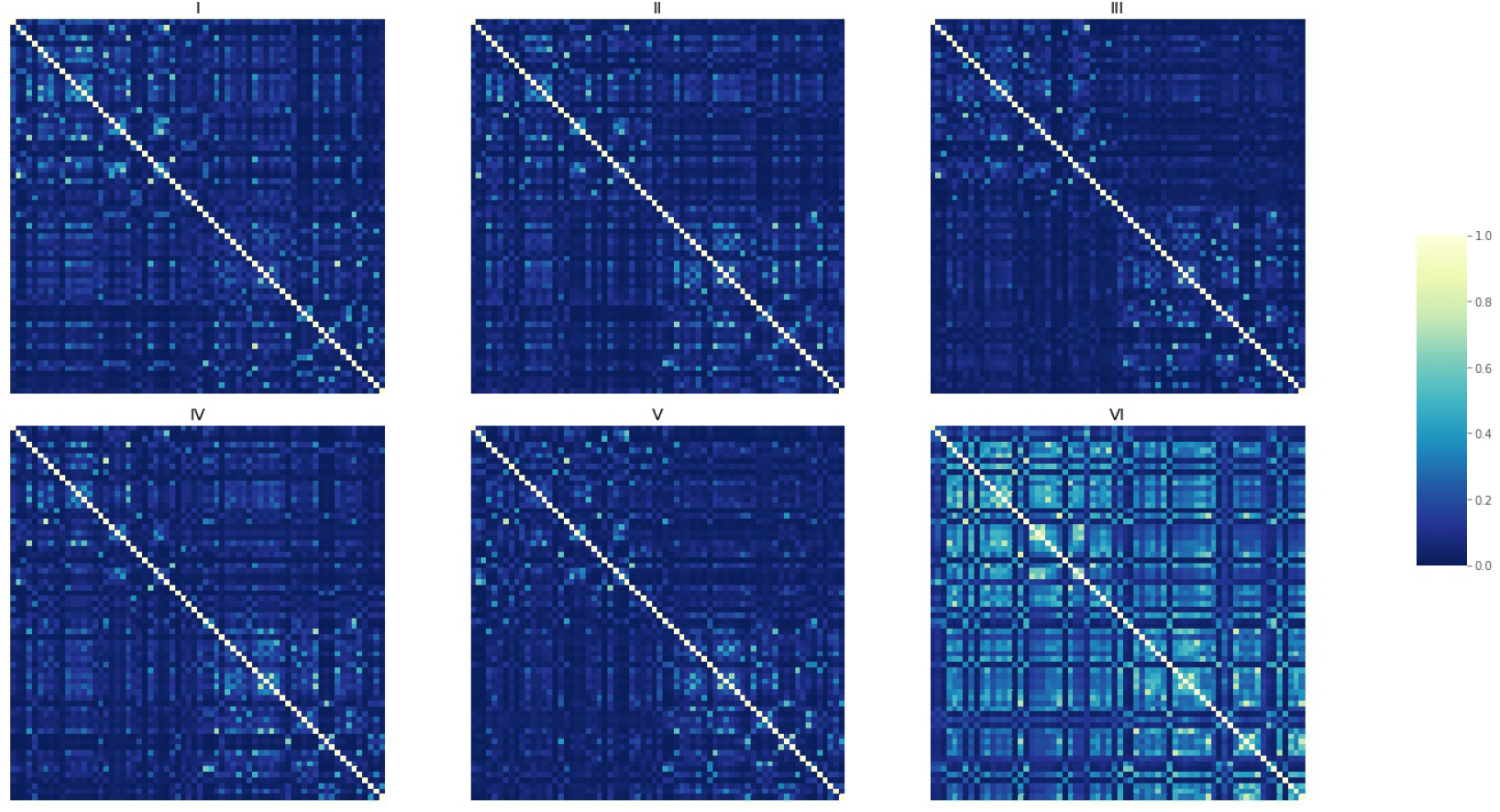
The pairwise connectivity over 68 ROIs from the second MEG resting state example.

**Figure 20:**
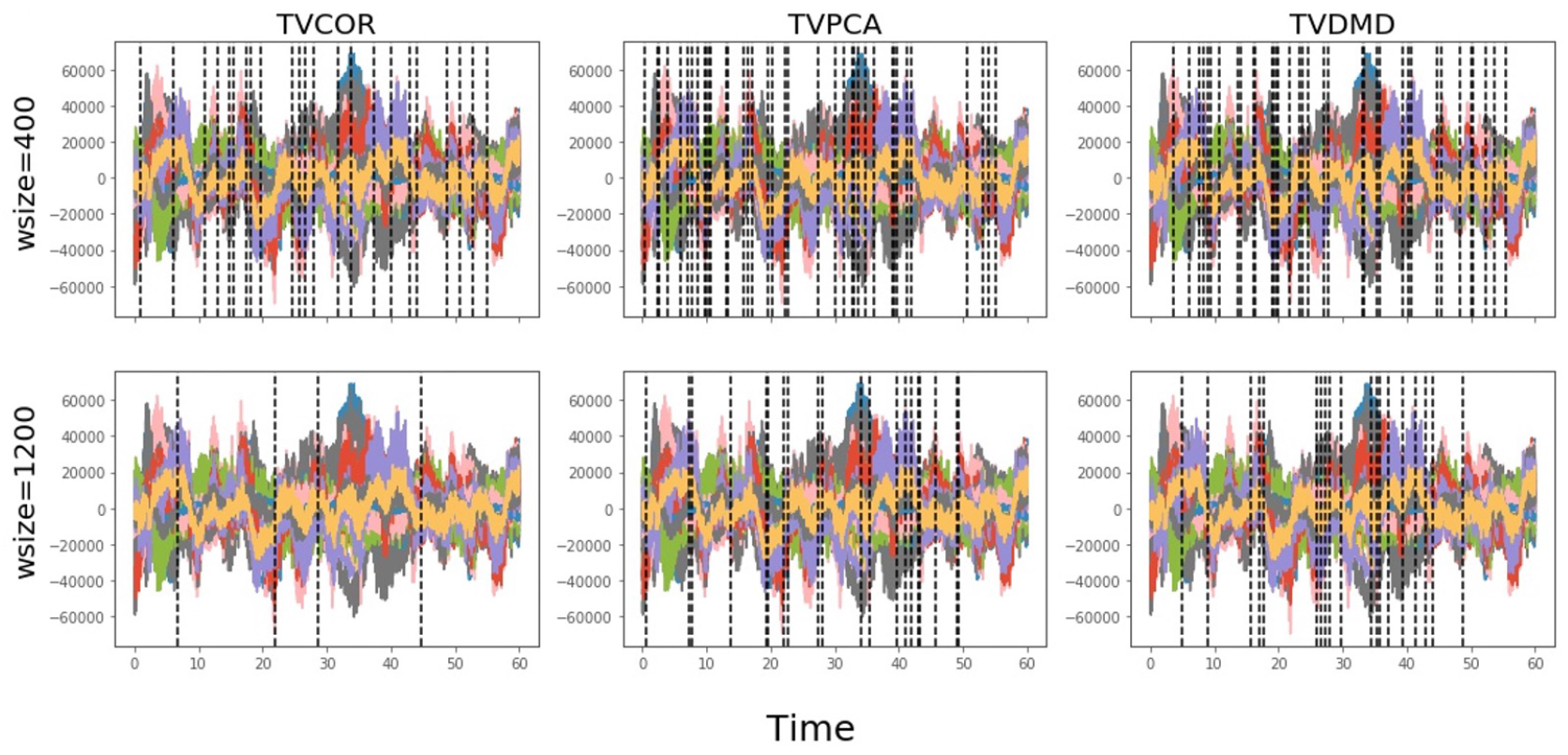
The brain state switch detection is not robust across different window size selections for the sliding-window approaches on the second MEG resting state example.

**Figure 21:**
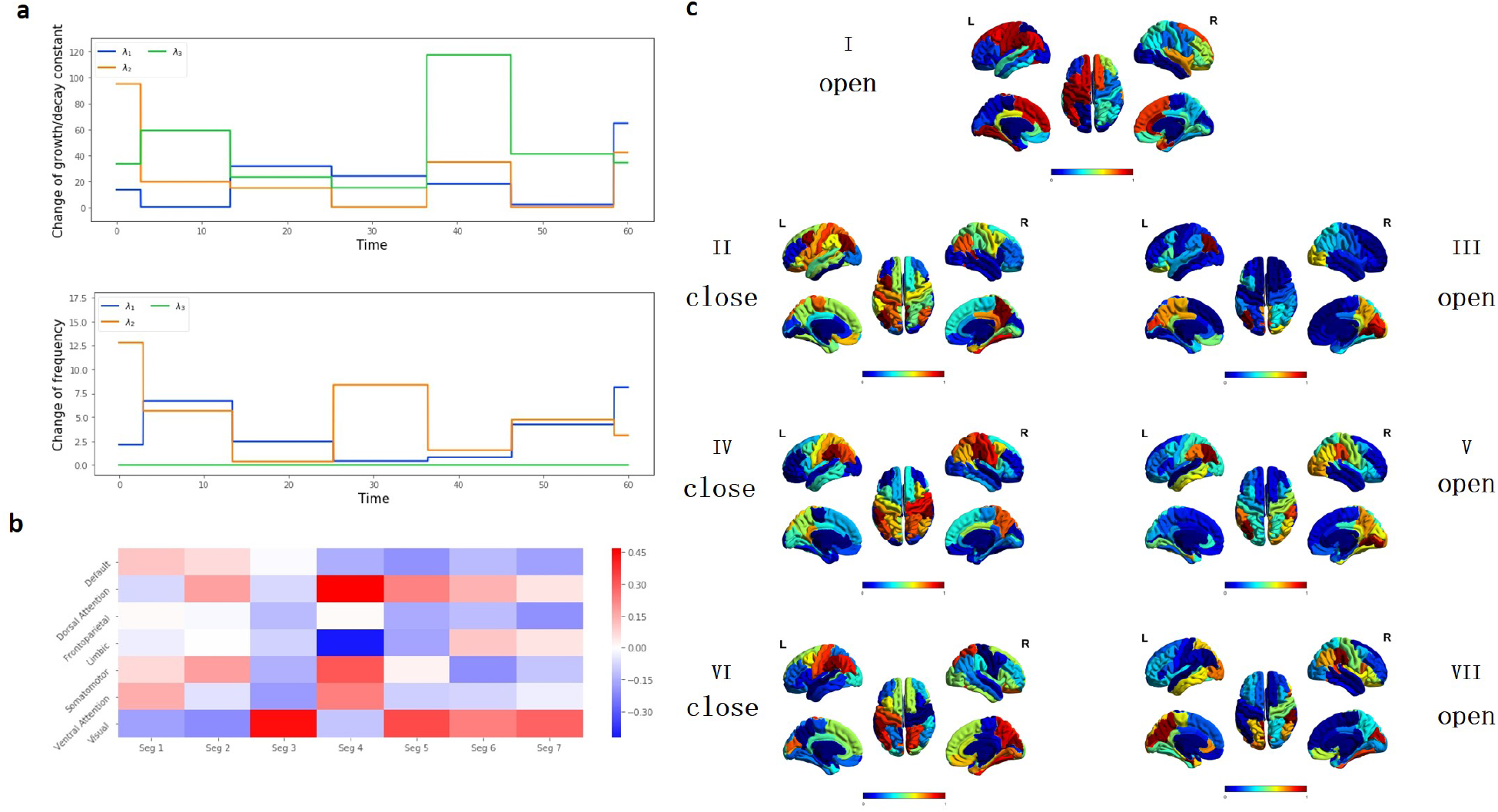
The results from the first eye-opening-closing MEG record. (a) Changes of growth/decay constant 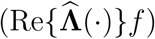, changes of the frequencies 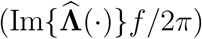. (b) The Pearson correlation between the weighted spatial features and the seven canonical networks. (c) The weighted spatial features across different segments detected by TVDN with eyeopening-closing labels.

**Figure 22:**
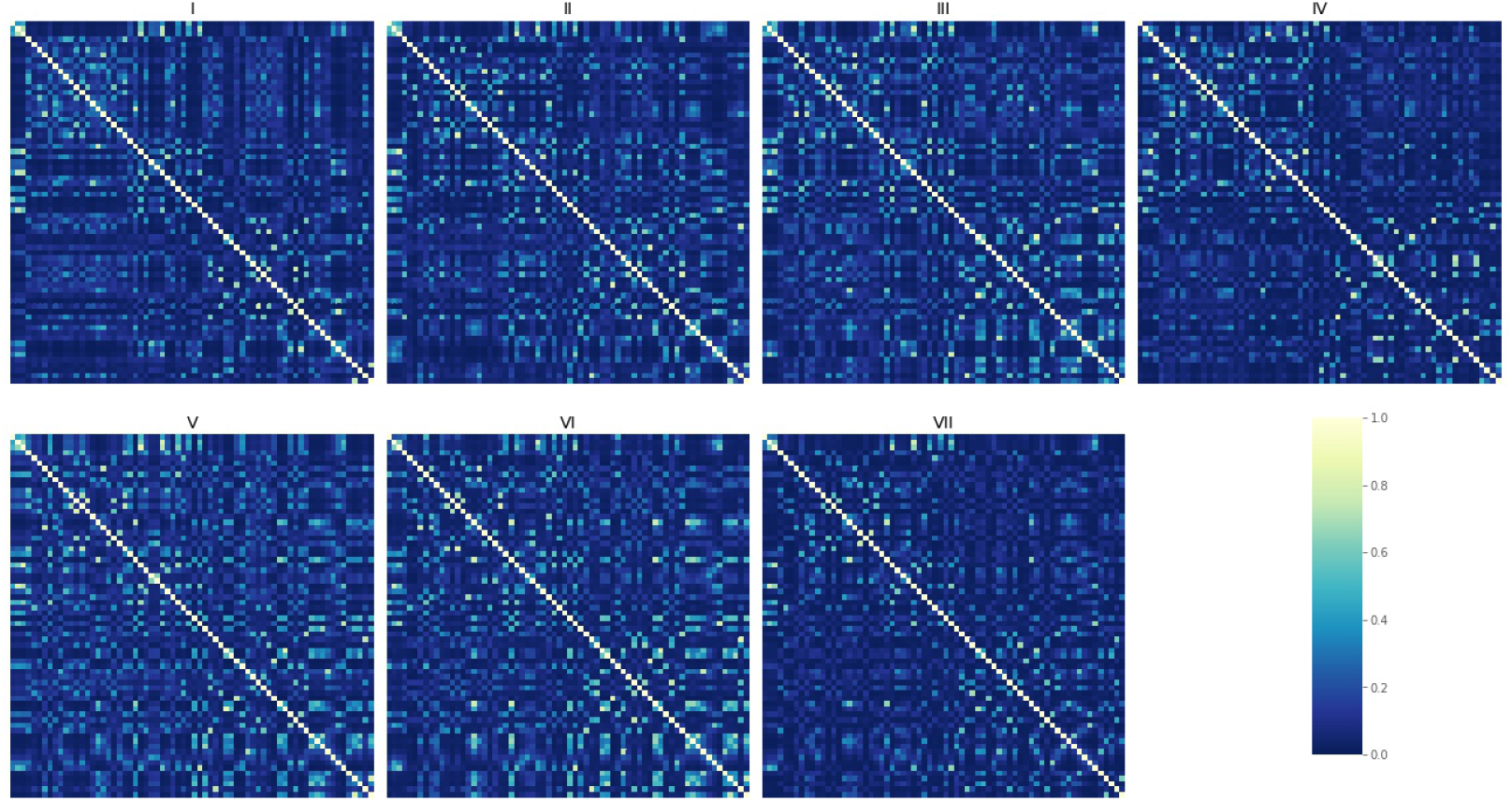
The pairwise connectivity over 68 ROIs from the first MEG eyes-open and eyes-close example.

**Figure 23:**
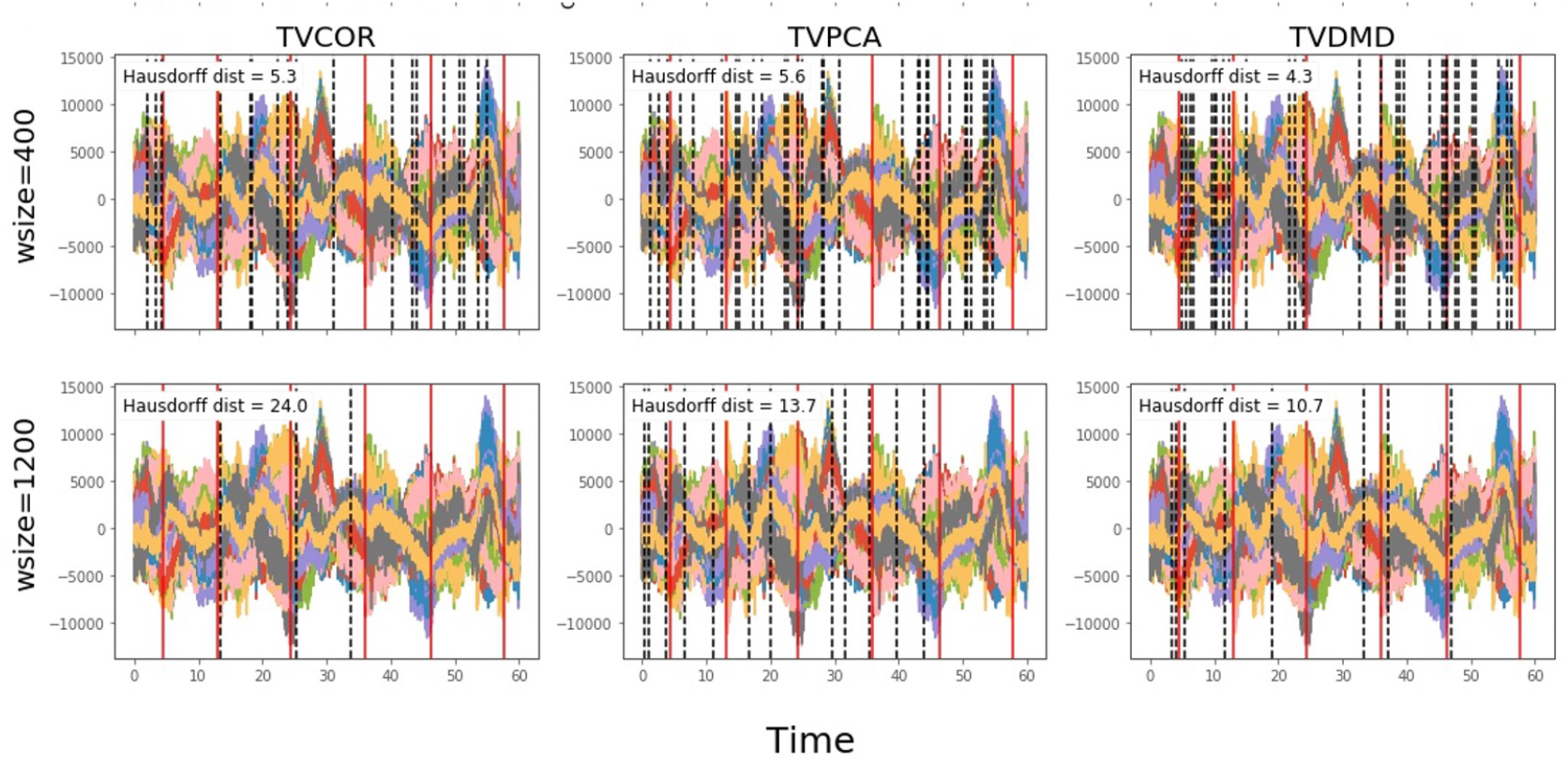
The brain state switch detection is not robust across different window size selections for the sliding-window approaches on the first MEG eye-opening-closing example.

**Figure 24:**
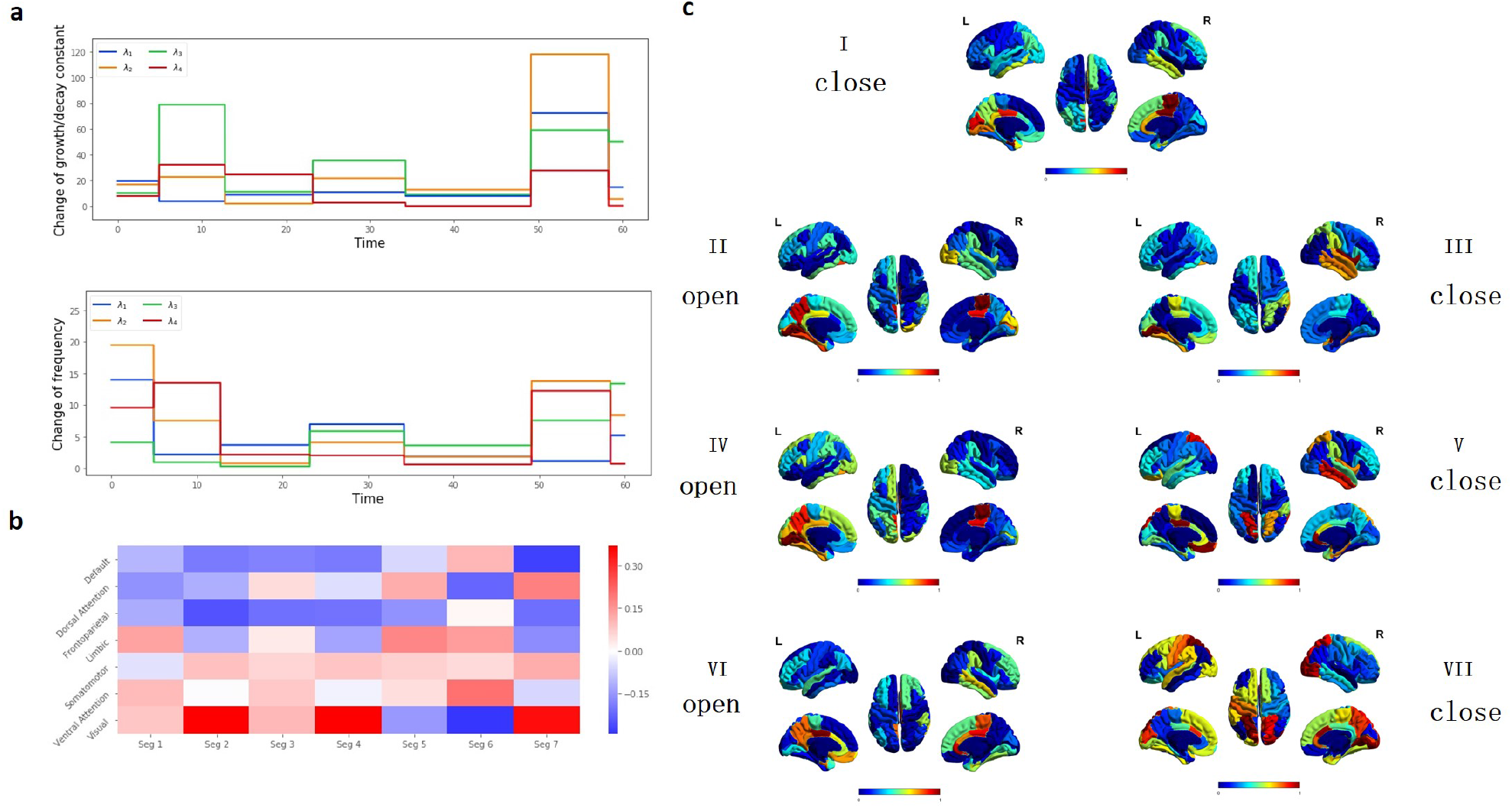
The results from the second eye-opening-closing MEG record. (a) Changes of growth/decay constant 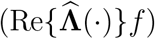, changes of the frequencies 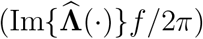. (b) The Pearson correlation between the weighted spatial features and the seven canonical networks. (c) The weighted spatial features across different segments detected by TVDN with eye-opening-closing labels.

**Figure 25:**
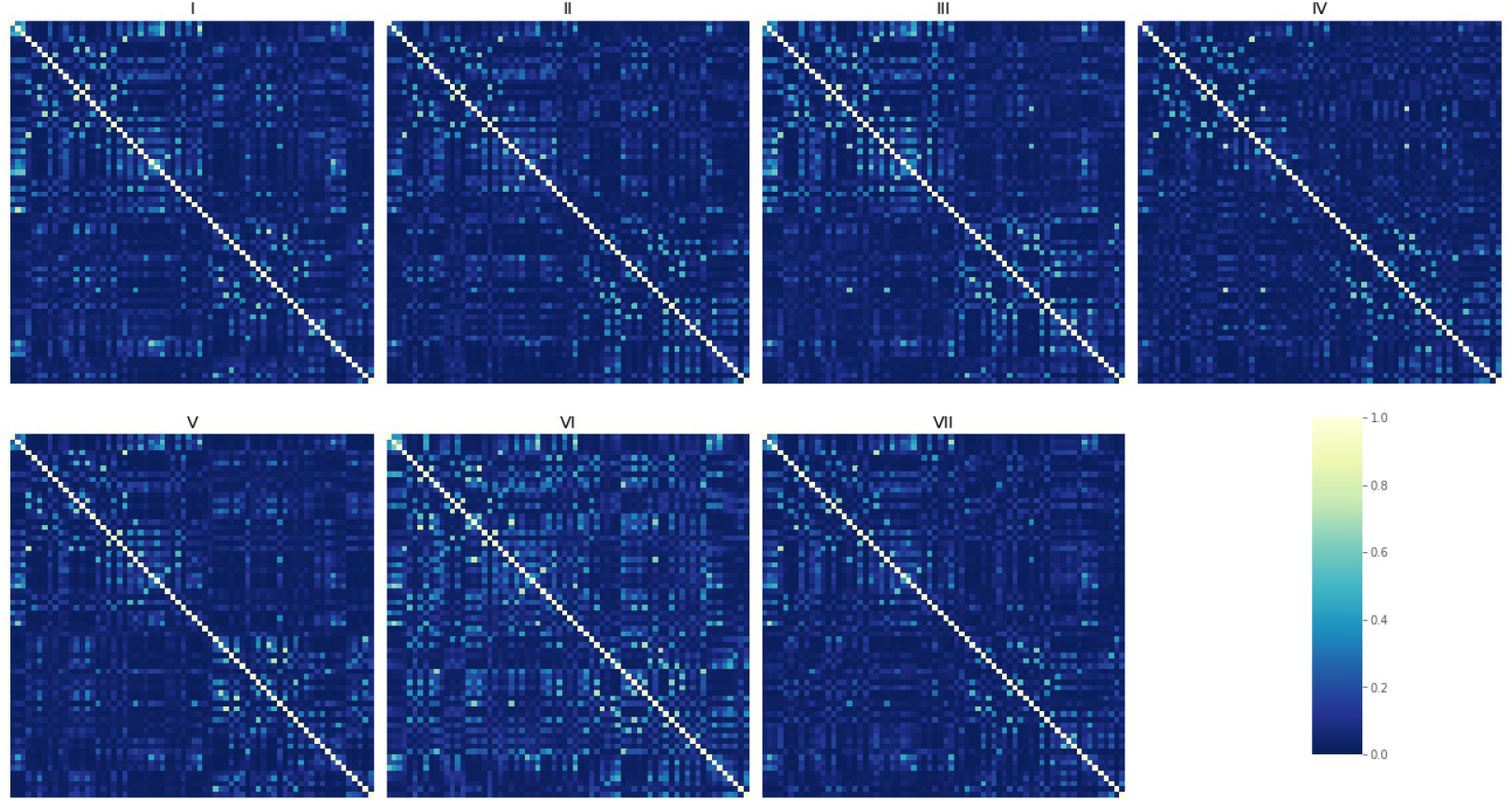
The pairwise connectivity over 68 ROIs from the second MEG eyes-open and eyes-close example

**Figure 26:**
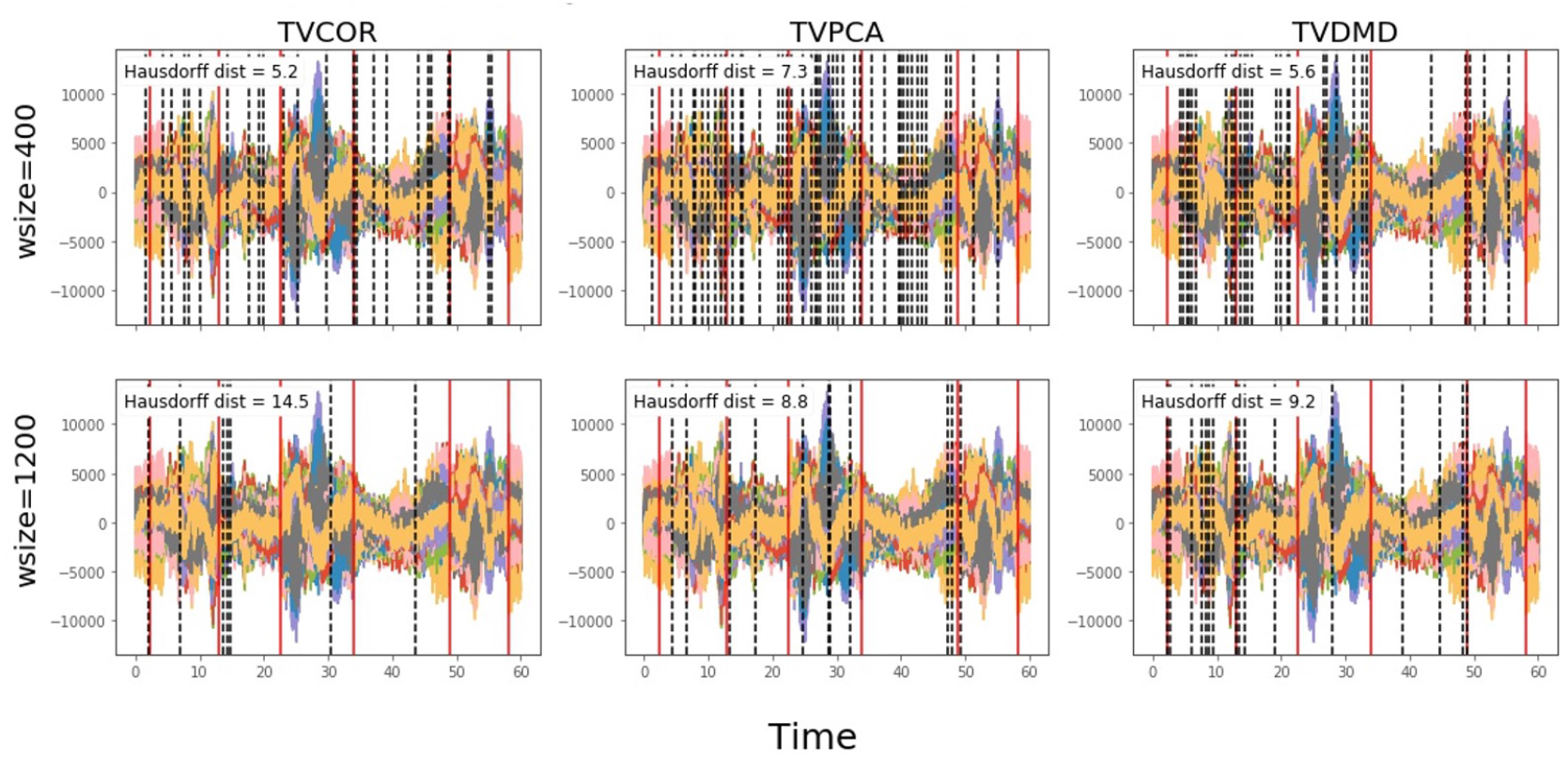
The brain state switch detection is not robust across different window size selections for the sliding-window approaches on the second MEG eye-opening-closing example.

